# A Bayesian modelling framework for inference of latent infection risk patterns from virus neutralisation assay titration data

**DOI:** 10.64898/2026.05.18.726027

**Authors:** Tarek A. Alrefae, Margarita Pons-Salort, Christl A. Donnelly, Ben Lambert, Everlyn Kamau

**Author notes:** joint senior authors (Ben Lambert, Everlyn Kamau).

## Abstract

Serological assays remain the standard approach for estimating the cumulative incidence of a pathogen and monitoring population immunity. Titration data from virus neutralisation assays are conventionally analysed using a nearly century-old interpolation-based method that neglects inherent imperfections in the assay and produces estimates with no measure of uncertainty. We introduce a two-part Bayesian framework for inferring latent infection risk directly from raw titration data. First, we develop a mechanistic model for serum antibody titration data that estimates antibody concentrations while quantifying uncertainty. Second, we propagate this uncertainty into an age-structured sero-catalytic mixture model by integrating over posterior draws of individual antibody concentrations, allowing joint inference on latent serostate membership, force of infection, and serological waning rate. We applied the framework to three cross-sectional serosurveys of the population of England, for enteroviruses A71 (EV-A71) and D68 (EV-D68) and coxsackievirus A6 (CVA6), which are leading causes of severe respiratory illness and hand, foot, and mouth disease. We estimated consistently higher and more persistent lifetime antibody concentrations for EV-D68 than for EV-A71 and CVA6. The proportion of recently infected individuals peaks around 40% by age 5 years for CVA6 and around 32% by age 7 years for EV-A71, before declining with age. These estimates are conditional on a model structure excluding complete seroreversion, which, where identifiable, implied a half-life of several decades. For EV-D68, the inferred proportion previously infected exceeded 50% by age 13 years and grew, with the force of infection declining more gradually with age. These estimates differ from those previously obtained from binarised versions of the same data. Our framework uncovers the wide-ranging variation in antibody levels that are often obscured by conventional endpoint titre methods and infers infection rates directly from raw titration data, without dichotomising at predetermined seropositivity cut-offs, while making minimal assumptions about virus-specific infection mechanisms.

**Author summary:** Serological tests measure antibody levels to estimate virus spread and protection from infection (immunity). Virus neutralisation assays dilute serum samples in steps and mix them with virus and cell cultures. The antibody titre is the highest dilution at which cells are protected from infection. Traditional analyses discard much of the testing information and overlook test imperfections. We present a two-part Bayesian framework that estimates antibody concentrations from the raw dilution data, carries the resulting uncertainty forward, and infers age-specific transmission of three common viruses – EV-A71, EV-D68, and CVA6. Our results show that EV-D68 infections may be more common, especially in children, than the other two viruses. This approach gives a clearer picture of seroconversion dynamics without dichotomising samples at fixed thresholds and has potential to improve estimation of virus transmission and population susceptibility.

## Introduction

Serological assays are essential for mapping pathogen-specific immunity landscapes and have long complemented symptom-based surveillance in infectious disease epidemiology [1]. The ability to detect antibodies makes serological assays useful for inferring population susceptibility, assessing herd immunity, and gauging vaccine-induced humoral protection [1, 2]. The virus neutralisation assay (VNA) is a widely used serological method that measures functional neutralising antibodies by evaluating specific interactions of serum antibodies and viruses. By measuring functional antibodies, VNAs enable more accurate assessment of antigenic variation in circulating virus strains compared to binding antibody assays, which detect all antibodies regardless of their neutralising (i.e. inhibitory) capacity. VNAs are inexpensive, require little specialist equipment, and follow standardised protocols, albeit are generally labour intensive [3].

In a VNA, a serum sample is serially diluted, mixed with a virus solution, and then applied to cultured cells. If neutralising antibodies are absent, the virus infects the cells, replicating and causing damage, otherwise known as a cytopathic effect (CPE). A CPE becomes visible after 3 − 4 days. When neutralising or protective antibodies are present and effective, they prevent a CPE (Fig. 1). The conventional output of a VNA is summarised as the endpoint neutralising antibody titre: the highest serum dilution that prevents a CPE. Surveillance studies interpret this value relative to seropositivity thresholds determined using positive (from known previously infected) and negative (from known previously uninfected) samples. In other cases, endpoint titres from paired serum samples are compared for incident seroconversion e.g., a ≥4-fold rise between paired serum samples is taken as evidence of a recent influenza infection [4]. While the endpoint titre indicates antibody function, it does not quantify the underlying antibody concentration. Moreover, VNAs typically test samples in a few dilution steps, often 8 − 10, which generates a limited number of discrete data points, limiting both the resolution of individual immune profiles and the depth of quantitative analyses at the population level.

**Figure 1:**
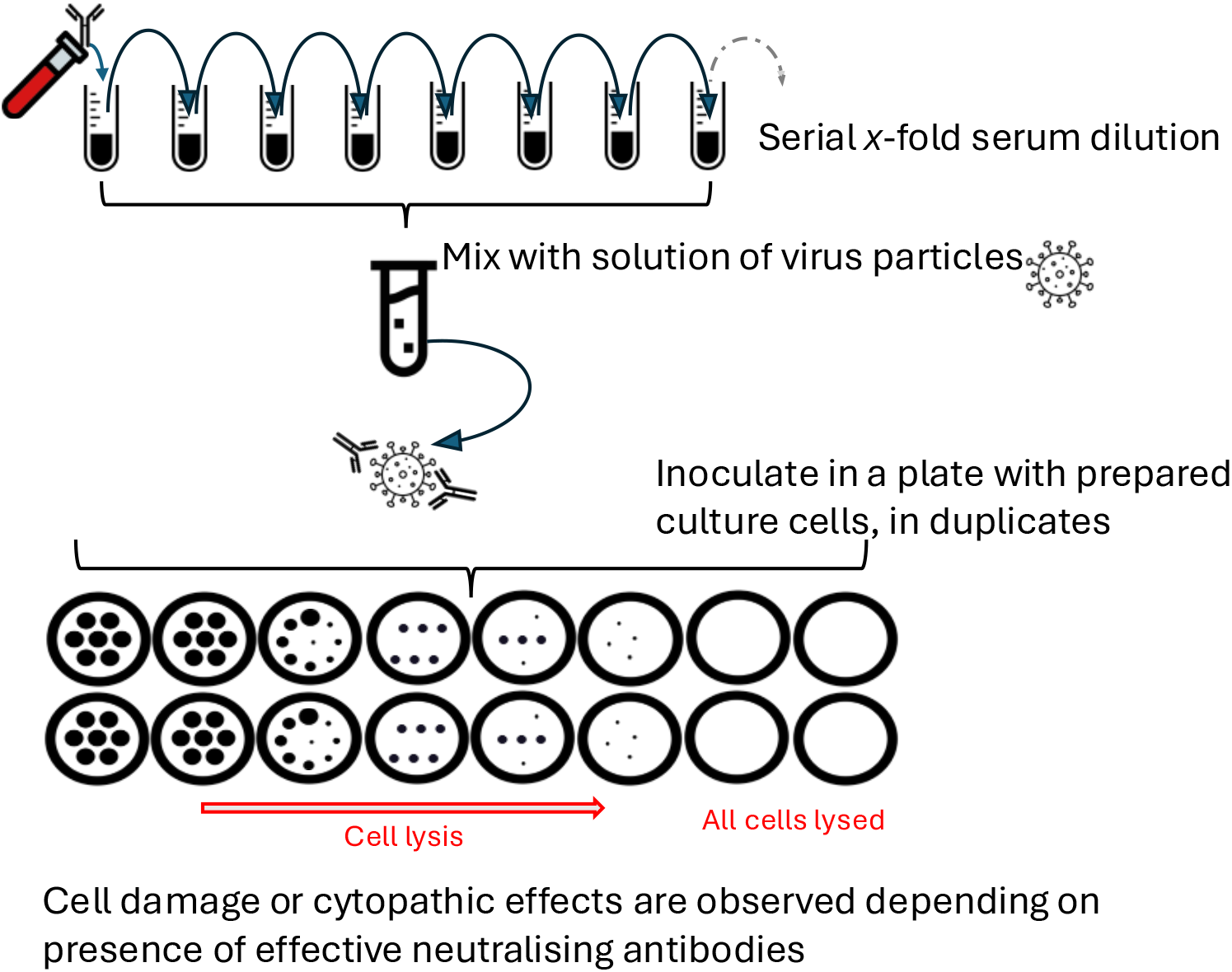
Virus neutralising assay. Illustration of neutralising antibody detection using virus neutralisation assays. Briefly, test serum is mixed with a virus solution and then inoculated onto cultured cells in a multi-well plate. In the absence of neutralising antibodies, live virus infects and lyses cells as it replicates. In the presence of sufficient neutralising antibodies, cell-virus interaction is blocked and lysis is inhibited. Cytopathic effects are typically assessed after 3-4 days.

We present a Bayesian analytical framework that uses VNA data to estimate the underlying serum antibody concentrations that are otherwise lost in the conventional endpoint summaries (Fig. 2). Related approaches have been developed for other settings. At the assay level, Bayesian hierarchical models have been used to estimate antibody titres from dilution-series data while correcting for batch-level experimental variation, for respiratory syncytial virus [5]. At the epidemiological level, serological models have treated serostatus as a latent variable to be inferred alongside transmission, for Mayaro virus in French Guiana [6], for West Nile and Usutu viruses in zoo animals in France [7]; in malaria serology, age-dependent antibody distributions have been modelled explicitly and the resulting uncertainty propagated forward into a reversible catalytic model [8, 9].

**Figure 2:**
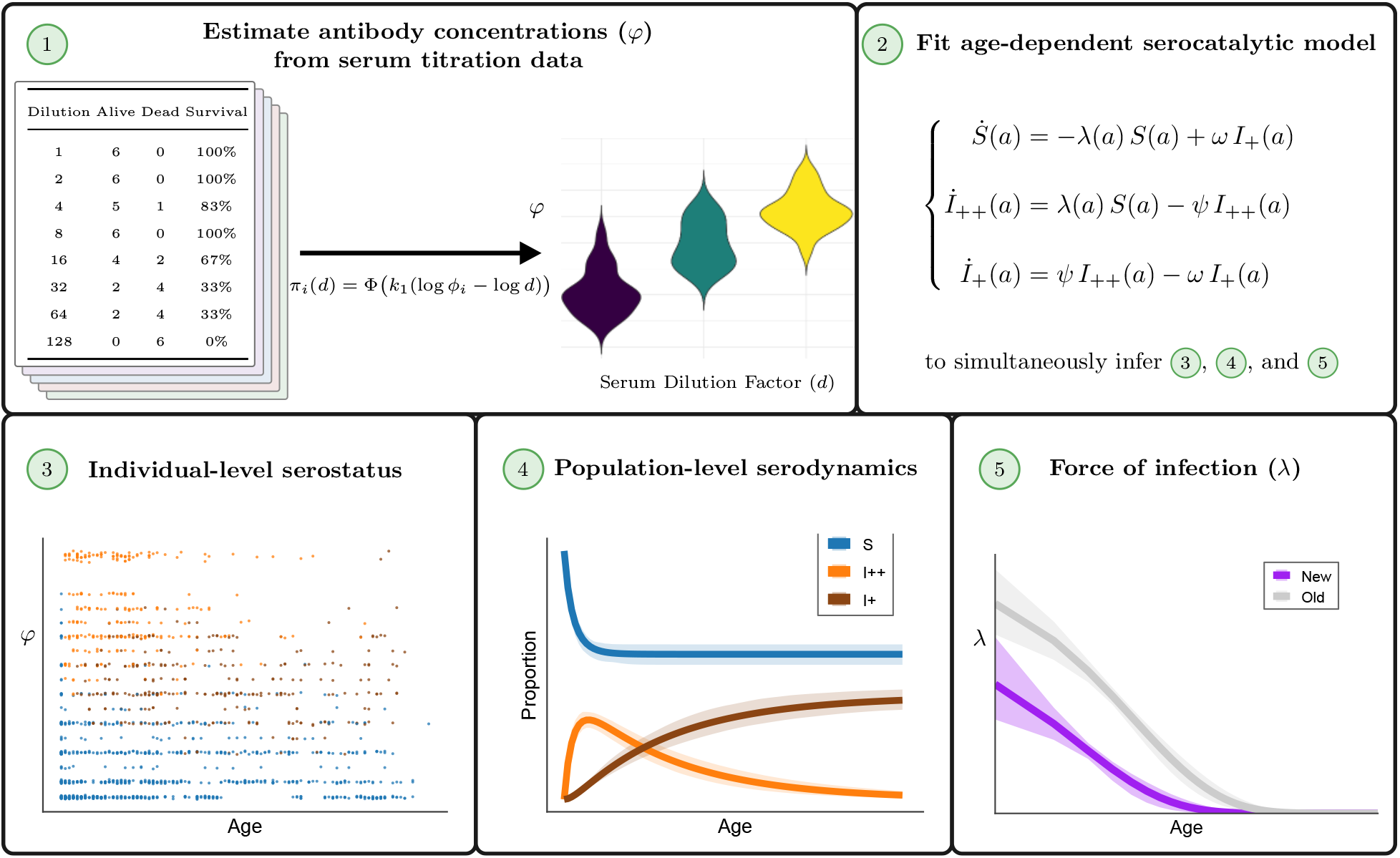
Inferring serological dynamics using virus neutralisation assay data: our methodological workflow. Briefly, serum titration data are fitted to a probit model to estimate latent antibody concentrations (*ϕ*) for each individual sample. Posterior draws of individual antibody concentration are then propagated into an age-dependent catalytic model to infer infection and immunity dynamics and to estimate the force of infection; we consider the force of infection to be strictly age-dependent. The colours in panel 3 and panel 4 correspond with the model compartments in panel 2: S susceptible (blue), *I*_++_ recently infected (orange), *I*_+_ infected a while ago (brown). Panel 5 illustrates comparison of a previous model fitting analysis (“Old”; where individual data were binarised as seropositive or seronegative based on a predetermined threshold) [23] with the present analyses (“New”; using the SII model) for EV-A71.

Our framework bridges the gap between these two largely independent strands. Assay-level models of dilution-series data have produced per-sample posterior distributions for antibody titres, but have then collapsed them to a point estimate for downstream use. Epidemiological models of serological data have propagated uncertainty in latent serostatus into downstream inference, but have taken a single summary measurement per sample as the observation. We estimate antibody concentrations from the raw well-level outcomes across the entire dilution series, using a mechanistic dose-response model of neutralisation as a threshold response to effective antibody exposure. We then couple the two stages by integrating over the first-stage posterior draws rather than fitting jointly, so that assay uncertainty enters the epidemiological likelihood while remaining separable (the assay model can be used on its own wherever titration data exist, independently of any epidemiological structure). Because a concentration rather than a serostatus is carried forward, the second stage can estimate not only whether an individual has been infected, but how recently.

We demonstrate the framework using simulated titration data and real cross-sectional VNA data from England, inferring the footprint of population immunity and age-dependent infection risk for three enteroviruses. These viruses are genetically and clinically distinct, they are major causes of respiratory illness and hand, foot, and mouth disease, and they occasionally lead to severe neurological or cardiopulmonary complications [10, 11, 12, 13]. The framework offers a principled means of estimating cumulative incidence and tracking immunity dynamics directly from serological data without relying on fixed or virus-specific seropositivity thresholds.

## Methods

### 1.1 Estimation of antibody concentration from serum titration data

#### A. Mechanistic motivation and statistical formulation

Virus neutralising antibody detection methods involve mixing solutions of known virus quantity with different serum dilutions, inoculating the mixture onto prepared cell cultures, and then evaluating the degree of cell damage or CPE as an indicator of presence (or absence) of effective neutralising antibodies (**Supporting Information** and **Fig. 1**). Usually, the serum dilutions are tested in duplicates or replicates to assess variability and enhance the reliability of output. In these assays, an antibody titre is usually determined as the highest serum dilution that inhibits CPE in >50% of replicate inoculations [14]. Suppose that for a given serum sample there exists (i) a true underlying concentration of neutralising antibodies, and (ii) a dose-response curve that governs how CPE occurs as a function of antibody concentration. With increasing serum dilution level, the probability of virus inhibition diminishes and CPE is increasingly likely. Consider a virus neutralisation assay setup as illustrated in **Fig. 1**. We model the probability of cell survival at dilution factor d ≥ 1 as

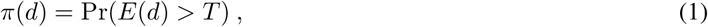

where *E*(*d*) > 0 denotes the effective exposure of the virus solution to neutralising antibodies and *T* > 0 is a threshold exposure, above which infection is prevented. The quantity *E* may be interpreted as an effective antibody count or volume. Because effective exposure may vary across orders of magnitude, we assume that relative to a reference exposure level *E*_ref_ > 0,

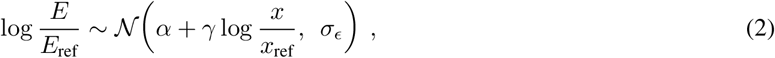

where *x* denotes the antibody concentration in the diluted serum solution, *x*_ref_ is a reference antibody concentration, and *α, γ*, and *σ*_ϵ_ are unknown constants. Writing the antibody concentration in the diluted serum solution as

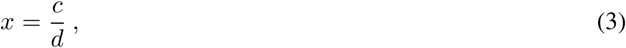

where *c* > 0 is the unknown underlying neutralising antibody concentration in the original serum sample, we obtain

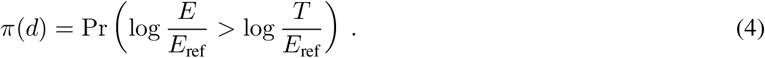

Standardising the normal random variable yields

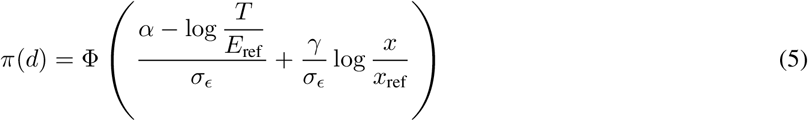

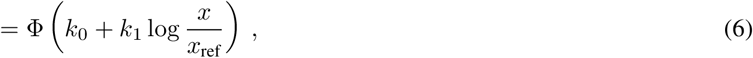

where Φ(·) denotes the cumulative distribution function of the standard normal distribution and

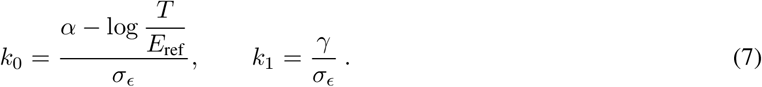

Substituting *x* = *c*/*d* gives

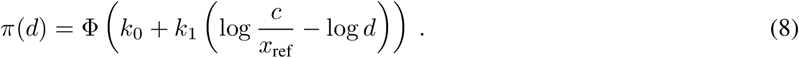

To simplify the parameterisation and improve identifiability, we choose the reference concentration *x*_ref_ such that *k*_0_ = 0 (see **Supporting Information** and **Fig. S1**). Under this choice, when *x* = *x*_ref_, the survival probability satisfies *π*(*d*) = Φ(0) = 0.5. Defining

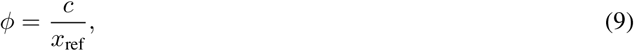

the dose-response model reduces to the probit form

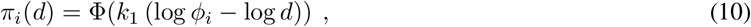

where *ϕ*_*i*_ > 0 is the latent antibody concentration parameter for individual *i* and *k*_1_ > 0 controls the steepness of the dose-response curve. In **Fig. S2**, we plot Eq. (10) for a range of hypothetical *ϕ* values, illustrating that as *ϕ* increases, cell cultures survive at higher serum dilutions. Supposing that the outcomes in each of the *n* dilution replicates are independent and identically distributed, then a binomial sampling model describes the aggregate number of cells surviving at a given dilution:

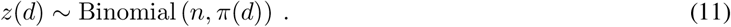

Equations (10) and (11) define the first-stage assay model, where Eq. (11) defines a likelihood function.

The model described above infers the dynamical process of neutralising antibody decay in assay measurements and we can fit this model to VNA data to generate joint posterior distribution estimates of *k*_1_ and *ϕ* in a Bayesian inference framework. The appropriate values of *k*_1_ depend on the units of *ϕ*. We place weakly informative half-Cauchy prior distributions on the slope and latent concentration parameters to allow substantial flexibility while retaining regularisation [15]. Posterior draws of log *ϕ*_*i*_ from this first-stage model are subsequently propagated into the second-stage serocatalytic mixture model described in Sections 1.2 and 1.3. We demonstrate implementation of the model using two serological case studies: (i) Reed and Muench’s hypothetical serum titration data, and (ii) cross-sectional enterovirus serology data from England.

#### B. Estimating (k_1_, *ϕ*) from a single serum sample: Reed and Muench’s hypothetical data

We assessed the practical suitability of the probit dose-response model in Eq. (10) using hypothetical titration data introduced by Reed and Muench in their classic study of endpoint titre determination [16]. Their example used hypothetical protective serum data from mice exposed to eight two-fold pathogen dilutions, ranging from 1:2 to 1:256, with six animals per dilution (**Table S1**). The Reed-Muench method uses a simple interpolation method without requiring specific computing software (see **Supporting Information** for a worked example).

Since six animals were tested at each dilution, *n* = 6 in Eq. (11) and the observed survival count *z*(*d*) can take any value in {0, 1, …, 6}. Given the substantial inherent randomness in such a small sample size, there can be considerable uncertainty of cell survival at each serum dilution. Eq. (11) effectively recasts the titration experiment as the outcome of a stochastic process and allows uncertainty in latent antibody concentration to be quantified probabilistically, rather than by interpolation alone.

We set the prior distributions of the probit model parameters *k*_1_ and *ϕ* as in **Table 1**. The model was run for 2000 Markov chain Monte Carlo (MCMC) iterations across each of 4 chains, with 1000 iterations discarded as warm-up using `Stan’s` default algorithm [17]. In **Fig. 3(A)**, we plot Reed and Muench’s hypothetical titration data (also presented in **Table S1**) and show the marked uncertainty in the survival estimates at each dilution (see also **Fig. S3**). The probit model fits the Reed and Muench titration data well (**Fig. 3(A)**) and their original estimate of the 50% endpoint dilution (26.9) is close to our Bayesian posterior estimate of the dimensionless initial undiluted antibody concentration, *ϕ* (median: 33.7). The two quantities are comparable by construction: setting *k*_0_ = 0 in Eq. (8) fixes the reference concentration *x*_ref_ such that *ϕ* is expressed on the scale of the 50% neutralising dilution; the Reed-Muench endpoint titre estimates the same underlying quantity (see **Supporting Information**). Our method has the advantage of also obtaining an uncertainty estimate, the central 95% credible interval (CrI): 17.8 – 70.6. Additionally, we can estimate the k_1_ parameter in Eq. (10), which characterises the shape of the dose-response relationship, as 0.69 (95% CrI: 0.40 – 1.05).

**Table 1:**
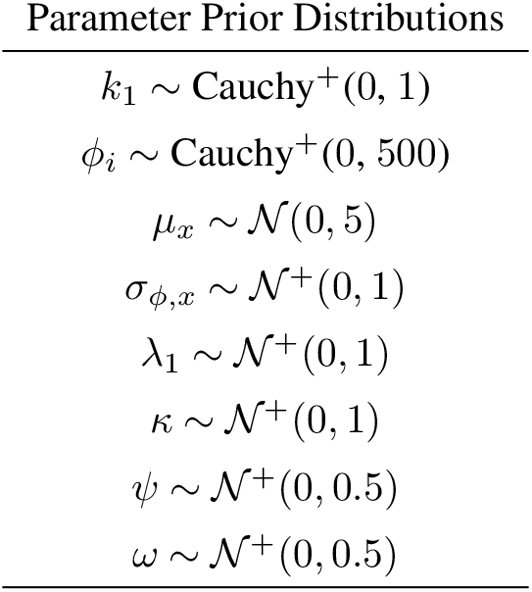
Prior distributions for the antibody concentration and serocatalytic mixture models. Here, *k*_1_ controls the steepness of the probit dose-response curve and *ϕ*_i_ denotes the latent antibody concentration for serosurveyed individual *i*. Cauchy^+^ and *N* ^+^ denote a zero-truncated Cauchy and normal distribution, respectively. For the three-state SII model, prior distributions were assigned as given above, with a single scale parameter *σ*_*ϕ*_ shared across serostates (this specification is used for CVA6 and EV-A71). For the two-state SI model fitted to EV-D68, the waning parameter (*ψ*) and the *I*_+_ location parameter (*µ*_*I*+_) were omitted and a separate scale parameter, *σ*_*ϕ*,*x*_ was estimated for each serostate. The seroreversion rate *ω* applies only to the SIS and SIIS models. Prior predictive checks for the probit antibody concentration model are demonstrated for EV-D68 in **Fig. S5**.

**Figure 3:**
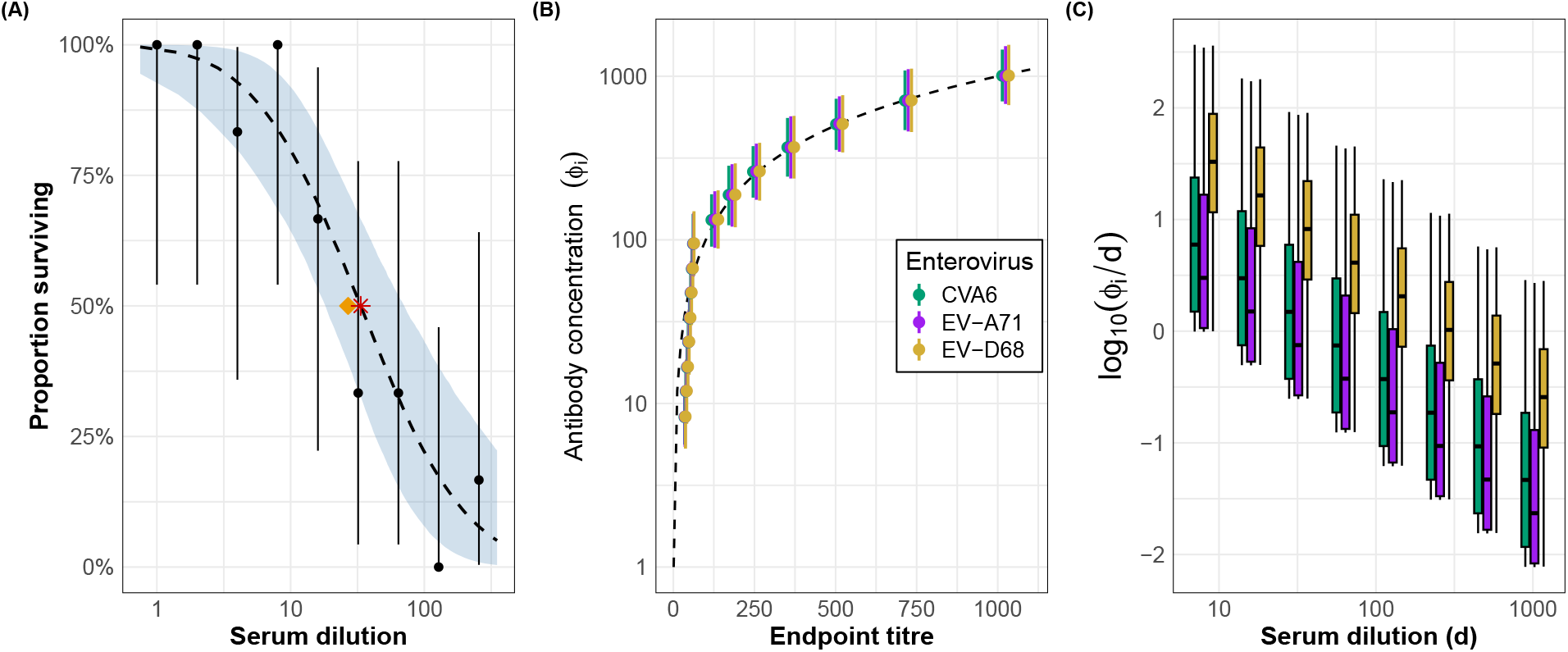
Output of probit model fitting to serum titration data. Panel (A) shows probit model outputs for fitting our statistical model to the Reed and Muench’s hypothetical titration data given in Table S1. The black points indicate the original data as reported in Table S1 and the error bars represent the 2.5 – 97.5% binomial proportion confidence intervals on the raw data. The blue ribbon represents Bayesian posterior uncertainty and shows the 2.5 – 97.5% modelled estimates of the proportion of surviving animals. The dashed black line represents the Bayesian posterior median survival, and the orange diamond indicates Reed and Muench’s original estimate of the 50% endpoint titre. The red star indicates the model-estimated 50% endpoint titre using the posterior median parameter values. Note that the horizontal axis is on a log_10_ scale. Panel (B) shows human serum neutralising antibody concentration against three enterovirus serotypes. This figure shows the median estimated serum neutralising antibody concentration (*ϕ*, vertical axis) as a function of endpoint dilution titres calculated via the Reed-Muench method (horizontal axis). The dashed black line is the identity, as setting *k*_0_ = 0 places *ϕ* on the scale of the 50% neutralising dilution. Antibody concentrations were estimated by fitting the mechanistic model in Eq. (10) to the serum titration data. Note that the vertical axis is on a log_10_ scale. Panel (C) plots the distribution of the log of the ratio between the model-estimated median antibody concentrations *ϕ* and the dilution level.

The statistical model in Eq. (10) is only identified up to the arbitrary reference scale used to define *ϕ*, and the prior distributions provide the regularisation needed for stable estimation. A consequence of this is that we get different estimates of the undiluted antibody concentration if we set different prior distributions, which requires setting the same prior distributions to ensure that concentrations are comparable across different experiments. This dilemma is not inherent to our model and is a consequence of inter-laboratory variability of serum titration outputs [18, 19]. Nonetheless, we demonstrate that the model can robustly handle serum titration data with noisy experimental output (e.g., dilutions 1:8 and 1:256 in **Table S1**).

To assess how far this dependence on the prior distributions propagates, we refitted the first-stage model under four alternative specifications, varying the scale of the half-Cauchy prior distribution on *ϕ* to 50 and 5000 (baseline: 500) and the scale of the half-Cauchy prior distribution on *k*_1_ to 0.25 and 4 (baseline: 1), as well as a replicate fit with the baseline prior distributions to establish a noise floor against which these contrasts could be read (**Fig. S4**). Estimates of *ϕ* were largely unchanged except under the wide *ϕ*-prior distribution, where the upper tail of the estimated antibody concentrations extended around an order of magnitude further. This reflects right-censoring rather than prior misspecification, as for individuals whose replicates survived inoculation at every measured dilution, the data bound *ϕ* only from below. Endpoint titres are subject to the same limitation, though they can only report a censored value (e.g. “>1024”) and offer no measure of uncertainty. The impact on serostate assignment was modest and symmetric about the baseline, and was concentrated among individuals whose assignment probabilities were already near 0.5. The effect on the second-stage force-of-infection estimates was negligible for all four alternative specifications.

#### C. Analysing enterovirus serology data

In practice, serum samples are analysed to monitor infection histories and immunity gaps in vaccination programmes. We assume that for actual serum samples with varying *ϕ*, the same dose-response function described above would govern the outcome. We apply our model to data from serum titrations against enterovirus D68 (EV-D68), enterovirus A71 (EV-A71), and coxsackievirus A6 (CVA6). Briefly, these are 1,588 serum samples collected as an approximately representative age-stratified cross-section of the population of England, in three surveys: 2006 (*n* = 517), 2011 (*n* = 504), and 2017 (*n* = 567). Participants ranged in age from 0 to 92 years (see **Supporting Information** and [20, 21, 22] for more details on these data). In the enterovirus serology data, each individual sample contains a set of paired dilutions and aggregate outcomes: 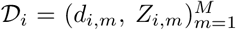, where *d*_*i,m*_ is the dilution factor (e.g., the “8” in 1:8) for an individual *i* in the dilution step *m*, and *M* is the total number of dilution steps evaluated in the titration assay. *Z*_*i,m*_ is the observed outcome, which is the number of replicates with surviving cell cultures in dilution step m. In the enterovirus serology datasets we analysed, *M* = 8, *d* ∈ {2^3^, 2^4^, …, 2^10^}, and *Z*_*i,m*_ can take any value from {0, 1, 2} because there were two replicates per sampled individual. **Fig. S5** shows the various unique experimental outcomes or patterns of surviving cell cultures at each serum dilution step and **Fig. S6** shows the serum titration data aggregated by age groups. We apply the dose-response function to the enterovirus serology data to estimate the underlying latent antibody concentrations (*ϕ*_1_, *ϕ*_2_, …, *ϕ*_*N*_) across all individuals (*D*_1_, *D*_2_, …, *D*_N_), where *N* is the number of serosurveyed individuals.

### 1.2 Modelling latent antibody concentrations

In this section, we develop serocatalytic models to describe how latent antibody concentrations evolve with age in response to infection. By modelling these dynamics, we can better distinguish stages of infection and classify individuals into various serostates based on their age and estimated antibody concentration levels. We draw a distinction between these inferred serostates and an individual’s serostatus, which is traditionally binarised into seronegative or seropositive. Before a primary infection, we assume there exists a small rate of antibody production resulting in a low equilibrium concentration of antibodies, *ϕ*_*S*_. Following infection, the rate of antibody production is temporarily elevated resulting in a higher equilibrium concentration, *ϕ*_++_, which then declines over time to a base rate resulting in an equilibrium concentration of antibodies, *ϕ*_+_. We assume that: *ϕ*_++_ > *ϕ*_+_ > *ϕ*_*S*_.

We assume that individuals at the time of sample collection are in one of three mutually exclusive serostates: susceptible (*S*), recently infected (*I*_++_), or infected a while ago (*I*_+_). We fit these models using ages and posterior draws of individual antibody concentration propagated from the first-stage assay model, in order to estimate the probability that each sampled individual belongs to a given serostate. **Fig. 2** provides a high-level overview of our methodological workflow.

### 1.3 Serocatalytic mixture models

#### 1.3.1 SII model

While the true underlying force of infection can reasonably be considered as a continuous function of both age and time, the cross-sectional nature of the available serological survey data and our desire for a simple, parsimonious model led us to only consider age dependence here. The serocatalytic model here is defined by a system of age-dependent ordinary differential equations (ODEs) to describe lifetime serological dynamics of a given birth cohort (*τ*). We assume the dynamics are determined by an age-dependent force of infection (*λ*) and a constant serological waning rate, *ψ*. We assume that antibody levels vary according to time of infection: before a first infection (e.g., after birth) individuals are considered susceptible to infection, *S*, and the measured antibody levels are typically low; after a recent infection, antibody levels spike and we model individuals as being in the *I*_++_ state; then over time, the detectable antibody levels wane and we model individuals as being in the *I*_+_ state. We refer to this as the “SII” model. The transitions between these serostates are governed by the following system of equations:

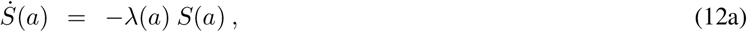

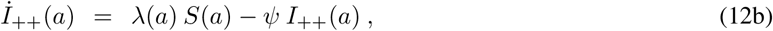

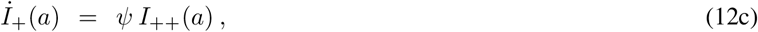

with initial conditions *S*(0) = 1 and *I*_++_(0) = *I*_+_(0) = 0. These initial conditions implicitly neglect potential passive immunity transferred at birth via maternal antibodies.

The SII model contains a serological transition *I*_++_ → *I*_+_ at rate *ψ*, in which antibody concentrations fall from a post-infection peak to a lower but still elevated equilibrium. It does not model complete seroreversion, *I*_+_ → *S*, in which an individual returns to the susceptible antibody distribution. We consider models incorporating seroreversion explicitly in Section 1.4, where we also review the empirical evidence: serological studies suggest that neutralising antibody detection for EV-A71 and CVA6 accumulates rapidly during childhood and remains common in adults, even though titres may decline after early adulthood [21, 23, 24].

Because the system of equations (12) is closed, these initial conditions ensure that *S*(*a*) + *I*_++_(*a*) + *I*_+_(*a*) = 1, for all ages *a* ≥ 0. Solving this system for a given individual *i* of a given age *a*_*i*_ yields a probability vector of the form {(ℙ (*i* ∈ *S* | *a*_*i*_), ℙ (*i* ∈ *I*_++_ | *a*_*i*_), ℙ (*i* ∈ *I*_+_ | *a*_*i*_)}, the elements of which are directly obtained from the model outputs.

Our analyses are also limited by the discrete, integer-valued ages recorded in the serological surveys. To accommodate this, we model the force of infection as piecewise-constant with year-length pieces [25]. That is, we assume that the force of infection 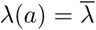 is fixed in the age range [*t* − *τ, t* − *τ* + Δ*t*], where t, *τ* are non-negative integers (Δ*t* must be a positive integer) and *t* ≥ *τ* . For example, taking Δ*t* to be a one year interval, an individual *i* born in 2020 (birth cohort *τ*_i_ = 2020) and sampled in 2025 is assumed to have experienced a sequence of forces of infection of the form 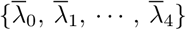 in their life up to the time of sample collection. Here, we assume this same set of forces of infection is experienced by any individual between birth and age 5 years, independent of their birth cohort *τ* . The governing equations in a given year within this age range become:

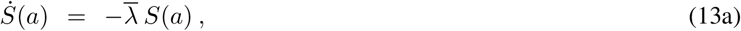

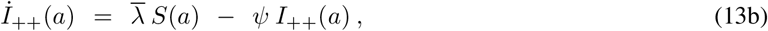

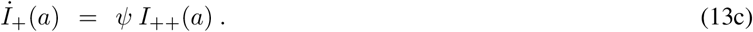

Eq. (13a) is a first-order separable ODE whose solution when Δ*t* = 1 year is:

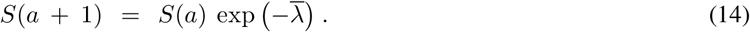

We can use an integrating factor approach to solve Eq. (13b):

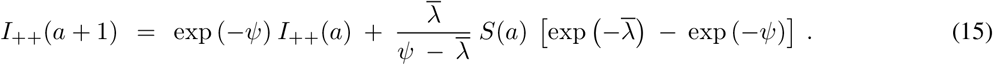

The set of solutions to system (13) is completed by setting:

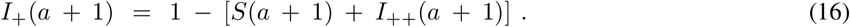

Equations (14), (15), and (16) are used to determine an iterative computational solution to system (12) under the assumption that the force of infection is structured in a yearly piecewise-constant fashion. Taking Δ*t* = 1 year yields the form we seek: between birth and some age at sample collection *a*_sample_, we model the total force of infection experienced by an individual of age *a*_sample_ as a sequence of forces of infection experienced for each age 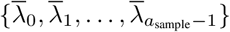. In practice, these one-year closed-form transition equations, Eqs. (14) – (16), were iterated over integer-valued ages to obtain age-specific serostate probabilities for each sampled individual.

#### 1.3.2 SI model

EV-D68 infections are highly frequent throughout a lifetime and studies have shown seroprevalence reaching nearly 100% by age 20 and remaining at that level throughout adulthood [22, 26, 27]. Given this and the robust immune responses typically elicited by EV-D68 [28, 29], we chose to use a simplified two-compartment model structure (“SI” model) for EV-D68 in contrast to the more detailed SII formulation above, which we used for EV-A71 and CVA6. In the SI model, individuals transition from being susceptible (S) to being seropositive (*I*_++_) and avoid any subsequent serological waning (i.e., *ψ* = 0 in equations (12b)/(12c) and (13b)/(13c)); it models the acquisition of the first infection only, despite the fact that infections are thought to occur frequently throughout a lifetime. The influence of maternal antibodies are still neglected by setting *S*(0) = 1 and *I*_++_(0) = 0 and we continue to exclude the possibility of seroreversion. The SI formulation was implemented as a separate two-state mixture model with latent classes corresponding to *S* and an absorbing infected state *I*_++_.

### 1.4 Seroreversion

To assess the consequences of excluding seroreversion completely, we additionally fitted models incorporating an *I*_++_ → *S* (two-state, “SIS”) or *I*_+_ → S (three-state, “SIIS”) transition at rate *ω*. The SIIS system is

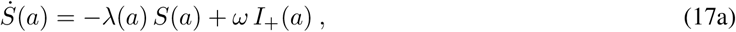

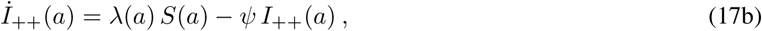

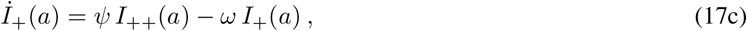

with the same initial conditions as the SII model (the SIS model is the two-state analogue of the SIIS model). Setting *ω* = 0 recovers the SII model exactly (the SI model is recovered by additionally setting *ψ* = 0, as before). We placed a half-normal prior distribution *ω* ∼ *N* ^+^ (0, 0.5) on the seroreversion rate, in line with that of the waning rate *ψ* (**Table 1**).

### 1.5 Model likelihood

For each individual *i*, the probit model fitting (Section 1.1) yields posterior draws

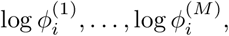

which we treat as a Monte Carlo representation of uncertainty in the latent antibody concentration *ϕ*_*i*_ . In the serocatalytic models (SI/SIS/SII/SIIS), we then integrate over these posterior draws when evaluating the state-specific likelihood contribution.

Let *X*_i_ denote the latent serostate of individual *i*, with *X*_*i*_ ∈ {*S, I*_++_, *I*_+_} for the SII model and *X*_i_ ∈ {*S, I*_++_} for the SI model. Conditional on serostate *x*, we assume

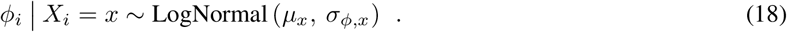

The scale parameters *σ*_*ϕ,x*_ may be estimated separately for each serostate or be constrained to a common shared *σ*_*ϕ*_. We compare both specifications for all three serotypes across all four candidate models in Section 1.6. A shared scale parameter proved a necessary identifiability constraint for the three-state models in most cases, and we adopt it for CVA6 and EV-A71. For EV-D68, the serostate-specific specification was clearly preferred and is used throughout.

Let *p* (*X*_*i*_ = *x* | *a*_*i*_) denote the age-specific probability of occupying serostate *x*, obtained from the solution of the serocatalytic model described in Section 1.3. For a given individual *i* and serostate *x*, we define the Monte Carlo approximation to the integrated state-specific likelihood

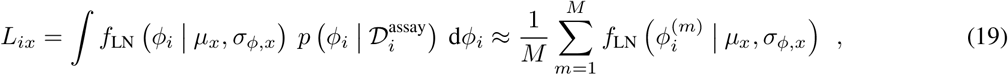

where f_LN_ (· | *µ*_x_, *σ*_*ϕ,x*_) denotes the lognormal density and 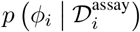 is the posterior distribution inferred by the first-stage antibody model fitting.

The likelihood contribution of individual *i* is then given by

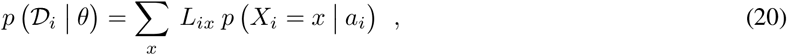

and the corresponding log-likelihood is l_i_ = log

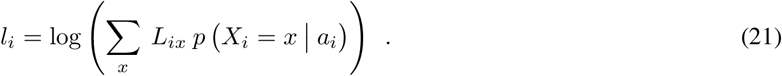

For numerical stability, all calculations were performed on the log scale using the log-sum-exp identity. Because the antibody posterior draws are available on the log scale, the lognormal density was evaluated as

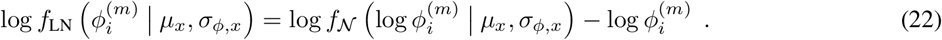

This formulation marginalises over both the latent serostate membership and uncertainty in the latent antibody concentration estimated from the assay model described in Section 1.1.

We constrain the serostate-specific location parameters *µ*_*x*_ to be strictly ordered such that *µ*_*I*++_ > *µ*_*I*+_ > *µ*_*S*_. Epidemiological knowledge suggests that exposure to and infection with enteroviruses including CVA6, EV-A71, and EV-D68 is highest in young children [30], and so we imposed the following structure on the age-dependent force of infection *λ*:

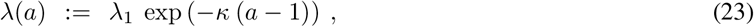

where *κ* > 0 and *λ* follows an exponentially declining relationship with age *a* ≥ 1 (our assumption of no transferred maternal antibodies means we do not consider potential prenatal infection risk). We had previously analysed the same enterovirus serology datasets by first determining seropositives and seronegatives based on predetermined seropositivity thresholds, and then estimating age-dependent force of infection using the same structure in Eq. (23) ([22, 23]). In the comparisons that follow we refer to the force of infection output from the previous analyses as “Old” and the output in this current work as “New”.

In the second-stage model, we estimate the serocatalytic and mixture distribution parameters *λ*_1_ and *κ*, the state-specific location parameters *µ*_*x*_, the serostate-specific scale parameters *σ*_*ϕ,x*_, and for the SII and SIIS models, the waning parameter *ψ*, and for the SIS and SIIS models, the seroreversion rate *ω*. Individual antibody concentrations are not re-estimated in the second stage; instead, uncertainty in *ϕ*_*i*_ is propagated from the posterior distributions generated by the probit model. **Table 1** shows the prior distributions for each parameter and our two-step fitting procedure is given in **Algorithms 1** and **2**.

In summary, the modelling framework involves two steps: (i) estimation of latent antibody concentrations *ϕ* from raw serum titration data by fitting the probit assay model given in Eq. (10) (Section 1.1); and (ii) fitting the serocatalytic mixture models by integrating over posterior draws of log *ϕ* propagated from the first step (Sections 1.3 – 1.5).

### 1.6 Model selection and convergence

We fitted all four compartmental structures (SI/SIS/SII/SIIS) with both shared and serostate-specific scale parameters, for each of the three serotypes. We assessed convergence in line with [31], namely 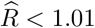 and bulk effective sample size (ESS) > 400 on all model parameters. A convergence summary for all 24 fits is given in **Table S2**.

Ten of twelve fits excluding seroreversion (SI/SII) converged, while only two of twelve fits including seroreversion (SIS/SIIS) converged with the seroreversion rate *ω* freely estimated. We observed that the failing SIS/SIIS fits were a result of confounding between the seroreversion rate and the force of infection parameters: this is a well-documented phenomenon for cross-sectional serological data [9, 25]. Models with serostate-specific scales converged for only four of twelve fits, while models with a shared scale parameter converged for eight of twelve fits. The failing three-state serostate-specific scale fits suffered from degenerate mixing, where *σ*_*ϕ,I*++_ collapsed towards zero around a small subset of individuals with high estimated antibody concentrations. The exception was the SII model for EV-A71, in which the *I*_++_ and *I*_+_ serostate location parameters were more well-separated (by 2.1 posterior standard deviations) than for e.g. CVA6 (1.4 posterior standard deviations).

Predictive accuracy was compared amongst models which converged by approximate leave-one-out cross-validation (LOO-CV) using the `loo` package [32] in `R` [33]. Results are given in **Table S3**. For EV-D68 the SI model with serostate-specific scale parameters was preferred by 27.1 (SE 7.5) and is used as our primary specification. For EV-A71 the SIIS model was preferred over the SII (both with shared scale parameter *σ*_*ϕ*_) by 8.9 (SE 4.4). We nonetheless retain the SII model with a shared serostate parameter as our primary specification for EV-A71, for reasons of comparability: we found the seroreversion rate *ω* to be unidentifiable for CVA6 under any specification, and adopting the SIIS model for EV-A71 alone would render any comparison between these two enteroviruses confounded by model structure. We report the SIIS findings as a sensitivity analysis (**Fig. S7, Table S4**) and quantify the consequences of excluding seroreversion in the **Discussion**. Readers whose interests lie in EV-A71 specifically (rather than in the comparison across serotypes) may prefer the SIIS estimates.

The mixture in Eq. (19) assigns a single location parameter *µ*_*x*_ to each serostate, pooled across ages, such that antibody concentrations are assumed to depend on age only through serostate membership probability, and not through the concentration distribution within a given serostate. To check this, we weighted each individual’s log *ϕ* by their posterior probability of serostate membership and evaluated the resulting age profile within each serostate, across six age bands ([0, 5), [5, 10), [10, 20), [20, 40), [40, 65), and 65+). We compared this against the range produced by data simulated under the fitted age-invariant model, for each serotype (**Fig. S8**). The assumption holds for CVA6 and EV-A71 in all serostates, and for EV-D68 in the recently infected state. The susceptible state for EV-D68 drifted upwards slightly with age more than the model predicts, which we consider in the **Discussion**.

### 1.7 Cut inference

Our two-stage inference procedure is a so-called “cut” approach [34]: information flows from the assay model to the serocatalytic model, but not back. We fitted a fully joint model for each serotype, in which the antibody concentrations *ϕ*_*i*_ are parameters and the serocatalytic mixture serves as their prior distribution. We found that all parameters across all serotypes were essentially unchanged, with the exception of *σ*_*ϕ,I*++_ for the EV-D68 SI model with serostate-specific scale parameters (cut: 0.83, 95% CrI: 0.73 – 0.94; joint: 1.13, 95% CrI: 1.04 – 1.22). This had no noticeable effect on the inferred transmission parameters (**Fig. S9**).

We retain our two-stage procedure as our primary analysis for reasons of interpretation. The two stages represent distinct mechanisms and their separation allows the first stage to be used on its own as a general method for estimating antibody concentrations with uncertainty from raw titration data. A fully joint model would condition these estimates on the particular epidemiological structures we considered. As presented, the first stage can be implemented independently of any subsequent epidemiological model.

We performed a simulation parameter recovery study using the SII model to assess whether the framework could recover known antibody and transmission parameters from synthetically generated data. We simulated antibody dilution measurements from known, randomly drawn antibody concentrations and analysed these with the probit dose-response model to evaluate recovery of *ϕ*. We then simulated a full age-structured serological dataset using known force of infection, waning, and serostate-specific antibody distribution parameters and fitted the complete two-stage inference pipeline. First-stage estimates recovered the individual *ϕ*_*i*_ across their full range, and the second-stage posterior distributions recovered the generating serodynamics and force of infection trajectories (**Fig. S10**).

### 1.8 Data and code

The Bayesian models were coded in `Stan`, using the `targets` [35] and `stantargets` [36] pipelines in `R`. For the probit antibody model, we ran four chains in parallel for 1000 iterations each, discarding the first 500 as warm-up iterations. Posterior draws of log *ϕ*_*i*_ from this first-stage model were thinned to 250 draws per individual before being passed to the second-stage model to control computational cost while preserving posterior uncertainty. The second-stage serocatalytic model was run with four chains in parallel for 200 warm-up iterations and 800 sampling iterations per chain. For both stages, convergence was assessed using the rank-normalised 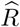 statistic and the bulk and tail ESS, with thresholds 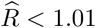 and ESS > 400 [31]. Second-stage models with a freely estimated seroreversion parameter *ω* were run with 800 warm-up iterations, 800 sampling iterations per chain, with the target average proposal acceptance convergence probability raised to `adapt_delta` = 0.9 [37]. **Table S5** and **Fig. S11** provide summary statistics (including relevant MCMC convergence metrics) and posterior density plots, respectively, for each of the fitted serocatalytic model parameters by enterovirus serotype for the best-fitting model. All analyses were conducted in `R v4.5.2` and the code and data are available on `GitHub` [38]. This includes a pseudonymised version of the original serosurvey data containing serum ID, age in years, dilution levels (1:8 to 1:1024), survival outcomes (recorded as 0 or 1), and survey year (one of 2006, 2011, or 2017).

#### Algorithm 1

Pseudocode for estimating antibody concentrations*ϕ*.

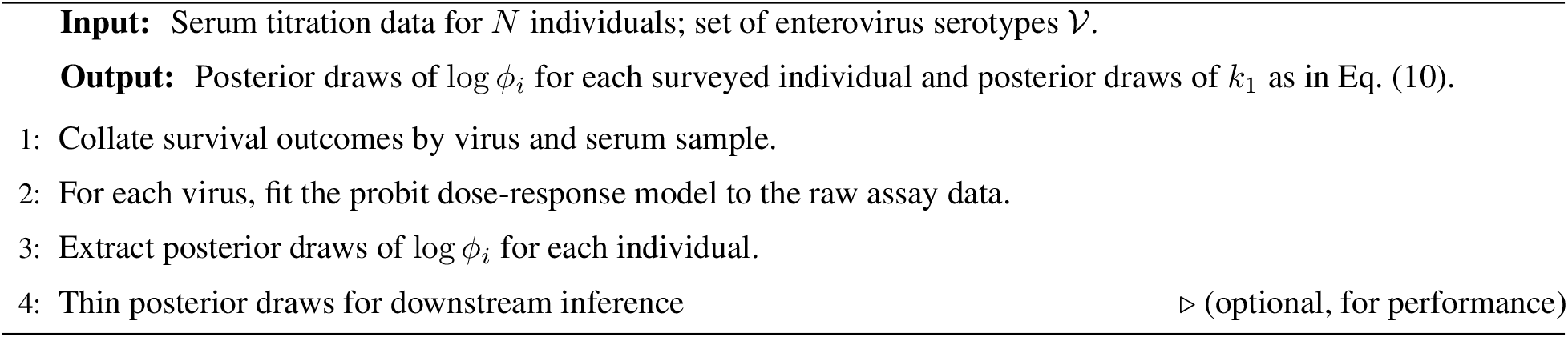

#### Algorithm 2

Pseudocode for estimating age-dependent force of infection λ(*a*) and serological waning rate *ψ*

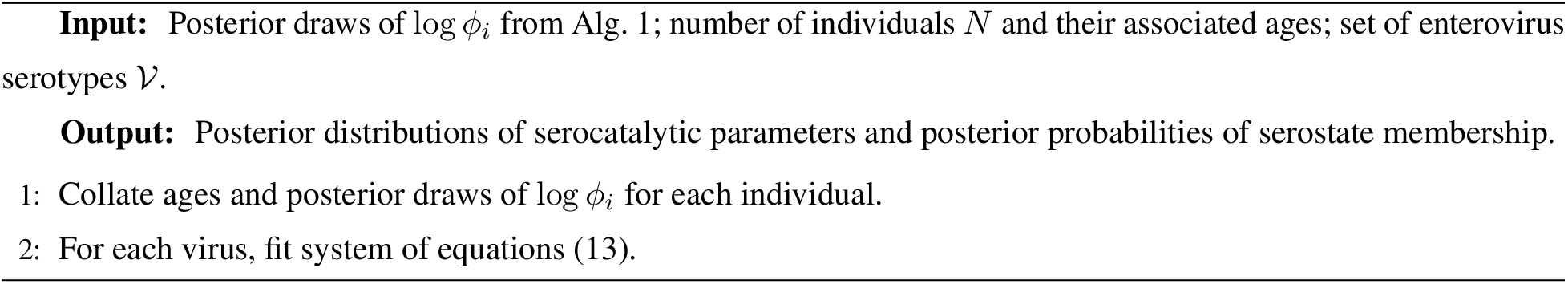

## Results

We estimated neutralising antibody concentrations (*ϕ*) for three enterovirus serotypes and, based on these estimates, inferred infection dynamics and immunity profiles across age groups. We describe our results in two parts: (i) estimation of antibody concentration from VNA data, and (ii) inference of serodynamics and force of infection from a serocatalytic model.

### Estimating antibody concentrations from VNA data

We fitted the probit model in Eq. (10) to datasets derived from serum neutralisation assays against EV-A71, EV-D68, and CVA6 to estimate neutralising antibody concentrations (*ϕ*) and the probability of cell survival at a given serum dilution. We estimated higher antibody concentrations against EV-D68 compared to EV-A71 and CVA6, with levels increasing gradually with age up to 80 years (**Fig. S12**). EV-A71 and CVA6 were comparable and showed more variability in the 5 − 35 year age range. The median individual-level *ϕ* estimates rose approximately 4-fold between birth and age 10 years for EV-A71 and CVA6, before gradually decreasing. As expected in a titration assay, for all three enteroviruses, antibody concentration declined monotonically with serum dilution (**Fig. 3(C)**).

We compared the estimated *ϕ* values with previously reported endpoint titres based on the Reed and Muench method [20, 21, 22] and found antibody concentration to be strongly positively correlated with the endpoint dilution titre for all three serotypes (**Fig. 3(B)**), confirming the suitability of using *ϕ* as a proxy for neutralising activity. In addition, our Bayesian method for estimating the underlying antibody concentrations results in quantifiable uncertainty measurements, which are not available with traditional point estimate endpoint calculations like the Reed and Muench method.

### Modelling infection histories and serodynamics

We estimated individual-level antibody concentrations (*ϕ*) and serostate assignments as a function of age for all serotypes (**Fig. 4**). For EV-A71 and CVA6, young individuals were estimated to have higher *ϕ* values and were more likely to belong to the *I*_++_ (recently infected) serostate: among individuals under age 10 years, 19.4% and 28.7% respectively were assigned to *I*_++_, compared with 2.5% and 2.8% among those aged 40 years and over, respectively. In older age groups, individuals with more moderate antibody levels were more often assigned to the *I*_+_ serostate, corresponding to those infected a while ago (29.6% and 39.3% of those aged 40 years and over, respectively), though with more relative uncertainty (5.8% and 0% of individuals assigned to the *I*_+_ serostate were assigned with probability ≥ 0.95, respectively, compared to 57.2% and 45.0% for the susceptible S serostate and 10.0% and 0% for the recently infected *I*_++_ serostate, respectively).

**Figure 4:**
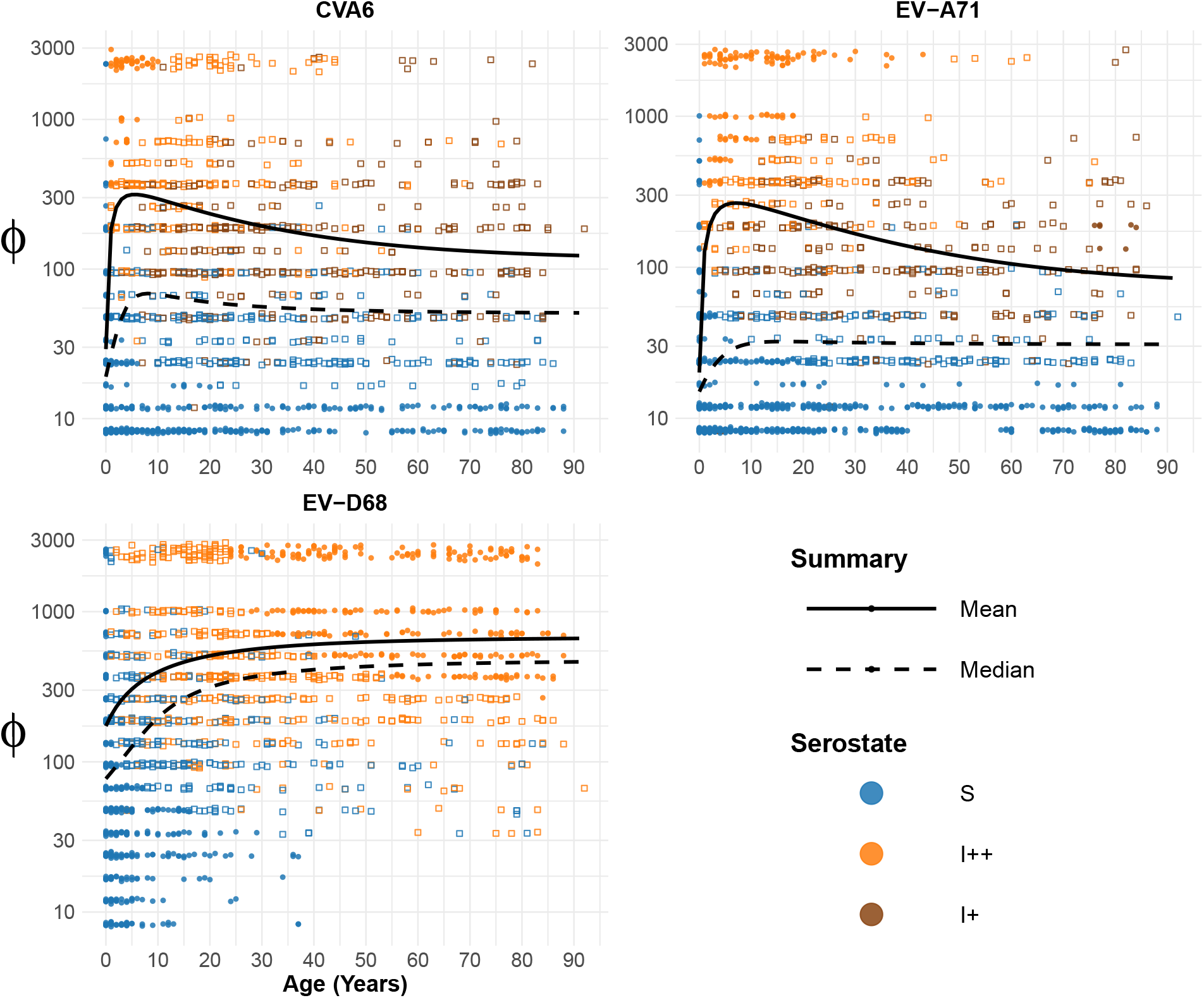
Inferred individual-level antibody concentrations and serostates. These plots show, for each enterovirus serotype, the estimated antibody concentration *ϕ*_*i*_ for each individual *i*, plotted against that individual’s age. Each data point represents individuals’ results for each specimen and is coloured by their model-predicted serostate (one of: susceptible (S, blue), recently infected (*I*_++_, orange) or infected a while ago (*I*_+_, brown)). The shape of the data points depict the posterior certainty of serostate assignment: solid dots represent at least 95% certainty of belonging to the respective serostate, while the open squares depict less certainty (< 95%) of serostate assignment. The solid (dashed) black lines represent the age-specific posterior mean (median) estimated antibody concentration across inferred serostate for the sampled population.

A further 68.0% (EV-A71) and 57.9% (CVA6) of individuals aged 40 years and over were assigned to the susceptible serostate, indicating appreciable residual susceptibility in the adult population for these two serotypes. For EV-D68, the observed dynamics were substantially different. We inferred higher *ϕ* values early in life, which remained elevated across age groups, indicating early and sustained exposure. Only 7.4% of individuals aged 40 years and over were assigned to the susceptible serostate, suggesting limited susceptibility in the adult population.

To summarise age-dependent changes in both antibody concentration and serostate composition, we calculated an age-specific, serostate-weighted mean *ϕ*, defined as the expected antibody concentration at each age after weighting the mean concentration of each serostate by the probability of occupying that serostate. Age-specific, serostate-weighted mean *ϕ* peaked at age 7 for EV-A71 (*ϕ* = 266) and at age 6 for CVA6 (*ϕ* = 315), followed by a slow decline. In contrast, the weighted mean *ϕ* for EV-D68 began to plateau from around age 30 at a substantially higher level (*ϕ* ≈ 600).

We reconstructed population-level serodynamics by plotting the proportion of individuals in each serostate as a function of age (**Fig. 5**). For EV-A71 and CVA6, the proportion of the population likely to be recently infected (I_++_) peaks between 30 − 40% around ages 7 and 5 years, respectively, before decaying. The *I*_+_ (infected a while ago) proportion increased with age, while the susceptible population plateaued between 50 − 60%. CVA6 showed faster decay in the *I*_++_ serostate proportion, reflecting a higher inferred serological waning rate; there was greater uncertainty across the inferred serostate proportions for CVA6 compared to EV-A71. EV-D68 showed considerably different serodynamics. The *I*_++_ proportion exceeded 50% by age 13 years and increased to near 90% by age 65 years. The bottom-right panel of **Fig. 5** shows the EV-D68 serodynamics fitted separately for each survey year (2006, 2011, and 2017). The inferred proportion susceptible was markedly higher in older individuals in the 2017 serosurvey than in 2006 or 2011.

**Figure 5:**
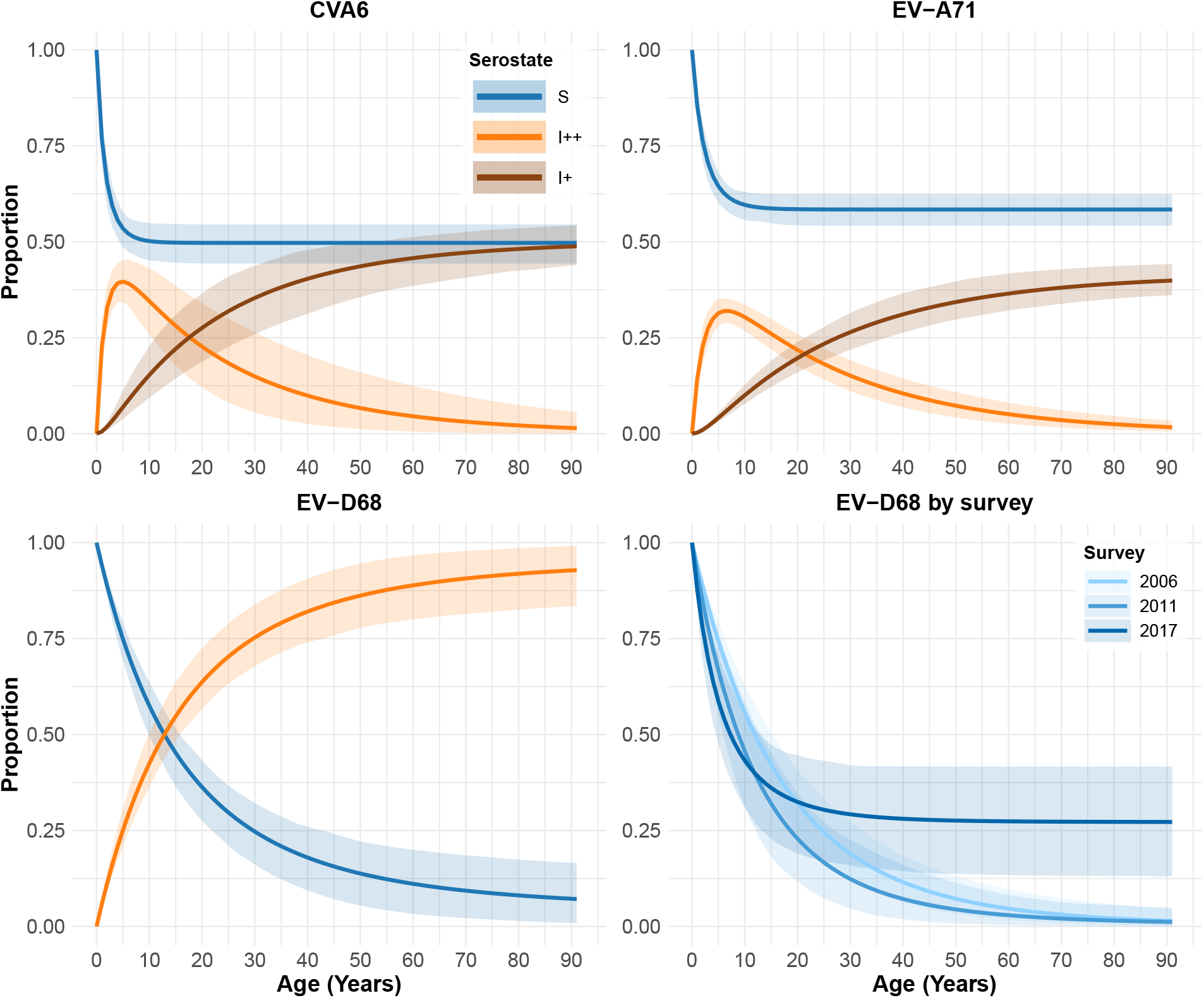
Inferred population-level serodynamics as a function of age for three enterovirus serotypes. These plots showcase the population-level age-dependent serodynamics corresponding to the individual-level modelled antibody concentrations and serostates in Figure 4. The modelled serostates include susceptible (*S*, blue) and recently infected (*I*_++_, orange) and in the case of CVA6 and EV-A71, infected a while ago (*I*_+_, brown). The solid lines depict median proportion of individuals in each serostate at each age. The shaded areas around each line depict the 95% central credible intervals. The bottom-right panel shows the EV-D68 susceptible serodynamics obtained by fitting the second-stage model separately to each of the three serosurveys (2006, 2011, and 2017), rather than to the pooled data (bottom-left).

Estimated age-dependent force of infection *λ*(*a*) is shown in **Fig. 6**. For CVA6 and EV-A71, the force of infection was relatively high at age 1 year and declined rapidly with age, approaching 0 by age 10 years. In contrast, the force of infection for EV-D68 started lower and declined only modestly with age. Performing the second-stage serocatalytic fitting separately by survey year shows that this pooled EV-D68 estimate averages over a time-inhomogeneous transmission regime (bottom-right panel of **Fig. 6**). The force of infection estimated using the 2011 survey was slightly higher compared to the 2006 survey, though both displayed comparable age dependence, maintaining low levels that only began to decline at relatively old ages. The force of infection estimated using the 2017 survey is noticeably different, beginning at a relatively elevated level at age 1 year and declining more rapidly with age: *λ*_1_ was estimated at 0.13 (95% CrI: 0.08 - 0.21) in 2017 compared to 0.06 (0.05 - 0.08) in 2006 and 0.08 (0.05 - 0.12) in 2011, marking an approximate doubling in the inferred force of infection at age 1 year between 2006 and 2017. Pooling therefore understates *λ*_1_ in the most recent survey. This change in the shape of the force of infection coincides with the 2014 global outbreak of EV-D68, which we discuss further in the next section. Stratifying CVA6 and EV-A71 by survey year yielded no comparable differences in the inferred force of infection (**Fig. S13**).

**Figure 6:**
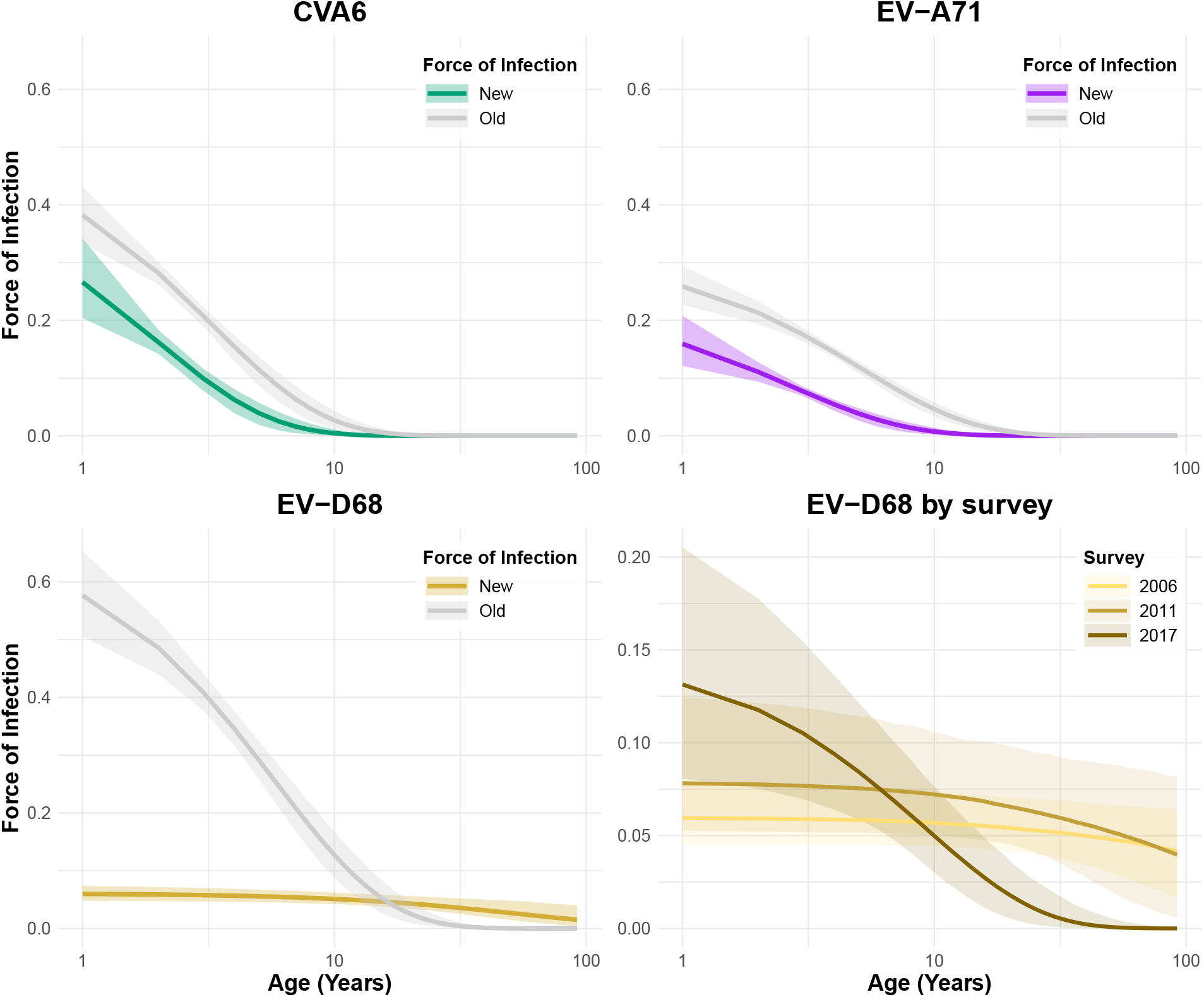
Inferred force of infection for three enterovirus serotypes. In each plot, the coloured line represents the median age-dependent force of infection *λ*(a) using the methods described in this paper (“New”, coloured line). These are compared to the force-of-infection estimates from previous methods (“Old”, gray line), as described in [23]. In the previous analyses, individual data were summarised as seropositive or seronegative (i.e., binarised) based on a predetermined threshold. The shaded areas in all panels represent the central 95% credible intervals in each case. Note that the horizontal axis is on a log_10_ scale. The bottom-right panel shows the EV-D68 force of infection obtained by fitting the second-stage model separately to each of the three serosurveys (2006, 2011, and 2017), rather than to the pooled data (bottom-left). Note the difference in scale of the vertical axis for the bottom-right plot.

We also compared the force-of-infection estimates obtained using our proposed two-stage Bayesian method (“New”) with previous estimates derived from a binarised version of the same underlying serological data (“Old”; see [22, 23]). These old estimates were obtained under a similar set of modelling assumptions, namely a purely age-dependent force of infection and excluding seroreversion (Model 5 in [23]); they are not a gold standard and this does not constitute a formal external validation of the framework. For CVA6 and EV-A71, both methods yielded similar age trends, but our new approach estimated lower force of infection values, especially in the 1 − 10 year age group, and in some cases produced narrower central 95% credible intervals in younger ages. For EV-D68, the new method suggested a lower force of infection across ages, in contrast to previous models that indicated meaningful age-structured variation. The new EV-D68 force-of-infection estimates also had lower uncertainty when pooling across the three serosurveys.

Two related features of the present approach plausibly account for our lower force-of-infection estimates. First, whereas our mixture model allows individuals with intermediate antibody concentrations to be apportioned probabilistically between serostates, older models binarised every individual above a certain threshold into a single seropositive class. Second, the SII model for CVA6 and EV-A71 provides a mechanism for age-dependent decline in antibody concentrations (*I*_++_ → *I*_+_, at rate *ψ*) that is not available to a binarised model. These earlier models instead must express the same age pattern through the force of infection alone. These combine to yield lower estimates of the force of infection *λ* in our model. Moreover, the available data cannot separate these two contributions, as the mixture likelihood and the *I*_+_ compartments are components of the same model and cannot be varied independently to compare against a binarised model.

### Seroreversion

The reported serodynamics are conditional on the assumption that individuals’ underlying antibody concentration levels, once infected, remain distinguishable from those individuals who remain susceptible throughout their life. We tested the validity of this assumption by fitting SIS and SIIS models to the data, to see if the inclusion of seroreversion might better explain the antibody concentrations inferred from the observed titration assay data. We found that ten of twelve SIS/SIIS fits across all three serotypes did not converge with a freely estimated seroreversion parameter *ω* (Section 1.6). Only the SIIS model for EV-A71 and the SIS model for EV-D68 converged.

For EV-A71, the inferred seroreversion rate was *ω* = 0.0128 per year (95% CrI: 0.0071 - 0.0199), corresponding to a seroreversion half-life of 54 years (95% CrI: 34.8 - 97.6). This estimate is compatible with two available empirical sources. In two prospective cohorts of children in southern China, naturally infected individuals were found to have a low probability of returning to a susceptible status [39]. A five-year follow-up study of EV-A71 vaccine cohorts recorded three seroreversions among 103 seropositive children over 532 person-years [40], implying a rate of 0.0056 per year (95% CI: 0.0012 - 0.0165). For EV-D68, the seroreversion rate in the converging SIS model was estimated to be *ω* = 0.00067 per year (95% CrI: 0.00002 - 0.00324), corresponding to a half-life of over 1000 years, effectively collapsing to the SI model.

For CVA6, where *ω* was not identifiable under any SIS/SIIS specification, we attempted a sensitivity analysis in which *ω* is held fixed at values corresponding to theoretically plausible seroreversion half-lives of 5, 10, 20, and 50 years (i.e. *ω* = 0.139, 0.069, 0.035, and 0.014, respectively). These fits also failed to converge, with 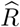 ranging from 1.15 to 2.16 across all tested *ω* values, against 1.001 to 1.004 for the SII model (where the seroreversion rate *ω* = 0). We therefore cannot report a seroreversion sensitivity analysis for CVA6.

We report the SIIS fit for EV-A71 in full in **Fig. S7** and **Table S4**. Allowing seroreversion yields a force of infection that declines more gradually with age and a susceptible proportion that exceeds the SII estimate from age 40 years onwards. The qualitative age pattern of the *I*_++_ serostate proportion is very similar in both models, peaking at 32.0% at age 7 years (SII) compared to around 33.6% at age 8 years (SIIS).

## Discussion

We developed and implemented a novel two-part Bayesian modelling framework where we first used a mechanistic probit model to estimate antibody concentration levels from VNA data before integrating these into a serocatalytic model to infer infection dynamics of enteroviruses EV-A71, EV-D68, and CVA6. Our approach yielded more nuanced and quantitative insights into the age-specific serodynamics and force of infection for these viruses compared to previous methods, without needing to impose an arbitrary seropositivity threshold.

### Methodological innovations

VNAs assess functionality of serum antibodies, i.e., detection of neutralising antibodies capable of preventing viral infection of cell cultures, or ability of sera to reduce CPEs. Neutralisation activity is measured through serum titration and the titration data are usually summarised as endpoint titres, which are then dichotomised into seropositive and seronegative values based on a predetermined seropositivity threshold for use as a proxy for past infection(s). However, this results in loss of information encoded, as endpoint titres provide semi-quantitative results but are not fit for accurate quantification of serum antibodies since the titration process decreases an unknown variable of antibody concentration by serum dilution. Also, the summary of serum titration data as endpoint titres obscures the range of antibody concentrations which could differ by several orders of magnitude. Here, we address this limitation by fitting a two-parameter probit function to raw VNA data to estimate serum antibody concentrations (*ϕ*). This probit formulation follows from modelling cell survival as a threshold response to effective neutralising antibody exposure, with serial dilution reducing the antibody concentration available in each well. It therefore represents the VNA as a dose-response experiment in which the probability of preventing CPE decreases as the serum dilution factor increases.

Our approach generalises beyond enteroviruses and is adaptable to any antibody titration-based system for which the SI or SII models are appropriate and where suitable VNA data exists. The method allows for robust quantification of seroconversion dynamics, can help improve downstream seroepidemiological modelling, and permits parameter inference without the need for predetermined seropositivity thresholds.

We also developed a Bayesian modelling framework that uses estimated antibody concentration (*ϕ*) values to probabilistically assign serostates to individuals. This allows for richer modelling of immune landscapes, beyond a coarse categorisation of individuals as either seropositive or -negative, and offers additional insights into recent and past infection histories. This modelling strategy enhances the interpretability of serological data and opens new avenues for quantitative seroepidemiology.

### Strengths and limitations

Our analyses are novel in that we demonstrate the ability to use a simple, mechanistically derived probit model to dissect antibody concentration from a conventional laboratory assay, the VNA. VNAs are the gold-standard tools for evaluating baseline serostatus and humoral responses to vaccination, and we now have a mechanistic way of obtaining quantitative data and more qualitative inferences from an assay that is pivotal in clinical medicine and laboratory science. We demonstrate the possibility of expanding simple and often overlooked laboratory output into a quantity amenable to Bayesian modelling. We have shown that serum antibody concentration estimates *ϕ* (i) mirror antibody titres, (ii) have the added benefit of associated quantified uncertainty (unlike conventional endpoint titres output by the Reed-Muench method), and (iii) are a suitable alternative data source for fitting age-dependent serocatalytic models.

Though the SII model provides a means for discerning between “recent” and “older” infections and allows for a deeper insight into latent infection risk patterns, it is a model that makes some key simplifying assumptions. The transition between serostates is modelled as strictly age-dependent, which neglects the inherently dynamic nature of immunity landscapes. This assumption is embodied in the functional form of Eq. (23), which requires that the risk of transmission be exponentially decreasing as a function of age. To test how this parametric functional form constrained our conclusions, we refitted each best-fit model with a piecewise-constant force of infection estimated independently over the age brackets [1, 2), [2, 5), [5, 10), [10, 20), and [20, 40] years (**Fig. S14**). For EV-A71 and CVA6 the age group piecewise brackets recovered the same qualitative pattern as the exponential functional form, where the force of infection was highest in the first years of life and declined steeply thereafter. For EV-D68 the piecewise estimates were likewise consistent with the flat age dependence obtained with the functional form. We conclude that the monotonicity constraint imposed by Eq. (23), though not driving the age patterns we report, does necessarily limit the resolution available in the first two years of life, where the force of infection changes most rapidly.

The assumptions we made regarding maternal immunity (i.e., how maternal antibodies may confer some level of protection against infection in newborns) are notable, though maternal immunity can be incorporated by adjusting the initial conditions. Maternally derived antibodies are expected to protect infants for roughly the first six months of life for EV-A71 [41]. Setting S(0) = 1 treats infants whose maternal antibodies have decayed as never having been protected, so that the apparent accumulation of infection over the first years of life is attributed entirely to primary infection, biasing *λ*_1_ (the inferred force of infection at age 1 year) upwards. This bias is expected to be largest for EV-D68, where the near-universal seroprevalence we infer in adults implies that a greater share of infants begin life with maternally derived neutralising antibodies [22, 27].

Excluding seroreversion has a predictable effect on our estimates. Without an *I*_+_ → *S* transition, any plateauing of seropositivity at older ages can only be accommodated by a force of infection that declines more steeply with age. This results in κ being biased upward, and the inferred adult susceptible proportion downward: comparing the SII and SIIS models for EV-A71 (where the seroreversion rate *ω* was identified), we find that median κ was 0.34 per year (SII; 95% CrI: 0.23 - 0.55) compared to 0.21 per year (SIIS; 95% CrI: 0.13 - 0.34) while the adult susceptible fraction at age 60 years was 58% (54% – 63%) against 67% (62% – 73%). Because we retain the SII model as our primary specification for EV-A71 (and the fact that seroreversion could not be identified for CVA6), the age-dependence of the force of infection we report for these two serotypes should be read as an upper bound (and consequently, the susceptible fraction as a lower bound). The qualitative pattern is unaffected, and the direction of bias is towards understating residual susceptibility in adulthood.

Our approach has computational costs relative to the Reed-Muench method, incurred by estimating the antibody concentrations *ϕ*, but these are likely to be relatively minor given modern computational resources, and are outweighed by the benefits of probabilistic inference. Here, the estimated antibody concentrations *ϕ* are dimensionless and lack formal laboratory units, which may limit their inter-laboratory comparability (this is by design, and one could conduct an analogous analysis with units if desired). However they remain meaningful within studies and provide a robust internal scale for analysis, with the added benefit of uncertainty quantification.

### Implications for enterovirus epidemiology

The inferred serodynamics and force of infection trajectories for EV-A71 and CVA6 align with established epidemiological patterns. For both of these enterovirus serotypes, we observed early-age infection, seroprevalence that peaks in childhood, and gradual waning of antibodies in adulthood. This closely mirrors the known epidemiology of hand, foot, and mouth disease and supports existing susceptibility estimates for these viruses [42]. In contrast, EV-D68 displayed dynamics which indicated early infection acquisition and persistent, high antibody concentrations into adulthood. This suggests frequent or repeated exposures over individuals’ lifetimes, consistent with previous studies that show widespread lifetime infection and reinfection [27, 43]. The observed shape of age-specific EV-D68 seroprevalence is similar to our previous analyses ([22]) and the overall dynamics of EV-A71 and CVA6 capture the observed seroprevalence dynamics in the original study ([21]). These demonstrate the ability of our methods and modelling approach to capture the true observed prevalence.

We inferred high susceptibility for EV-A71 and CVA6 – over 50% across all ages above 5 years – which suggests a risk of future outbreaks in the study population. This coincides with recent estimations of susceptibility to enteroviruses and is consistent with the potential utility in vaccinating young children against these enteroviruses [44].

The susceptible-state location parameter was estimated at *µ*_*S*_ = 4.35 (95% CrI: 4.21 – 4.48) for EV-D68, compared to 2.95 (2.85 – 3.04) for CVA6 and 2.71 (2.65 – 2.77) for EV-A71 (**Table S5**), suggesting that individuals assigned to the susceptible state for EV-D68 are assigned substantially higher baseline antibody concentrations than for the other two enterovirus serotypes. This elevated baseline was consistent across all three survey years (2006: 3.63 (95% CrI: 3.47 – 3.79); 2011: 4.60 (4.41 – 4.80); 2017: 4.43 (4.17 – 4.69)) and is potentially indicative of cross-reactive antibodies [45] or higher background seroreactivity to EV-D68 [46]. Another plausible explanation is the lack of an *I*_+_ compartment when modelling EV-D68 dynamics, which must therefore accommodate individuals with intermediate antibody concentrations within either the susceptible or recently infected distributions, potentially inflating *µ*_*S*_ and depressing the inferred force of infection. The age-stratified posterior predictive check in Section 1.6 is consistent with this: susceptible-state antibody concentrations for EV-D68 drift upwards with age slightly more than the fitted model predicts (though no such drift is present in the recently infected serostate for EV-D68). Distinguishing between these possibilities would require assay data from multiple co-circulating serotypes of related enteroviruses, which we do not have. The absolute value of the serostate location parameters should accordingly not be interpreted as a measure of baseline reactivity that is comparable between serotypes.

The apparent age-dependent force of infection for EV-D68, an enterovirus that causes respiratory illness, contrasts with the existing epidemiological knowledge of high levels of susceptibility and disease in those <10 years of age [28, 47]. The high potential for household transmission of respiratory viruses between children and adults could explain this finding [48]. A global outbreak of EV-D68 in 2014 ([49, 50]) and a sharp rise in reported respiratory disease associated with EV-D68 since 2009 ([51]), both of which lie within the period our data was collected (2006 − 2017), further complicate the analysis. Fitting the model separately to each survey (**Fig. 6**) shows that the pooled estimate averages over a changing transmission landscape. Though the apparently flat age dependence can be attributed to non-stationarity in the study period, we note that the most recent survey was conducted only three years after the 2014 global outbreak of EV-D68, meaning that the contrasts between this and the pre-outbreak surveys is likely to understate the true difference in transmission (as all individuals aged at least 4 years in the 2017 survey had lived through both pre- and post-outbreak periods). Time-varying transmission for these data has been estimated previously using a different, threshold-based approach [22]; we adopt the age-only framework here as a deliberate simplification to highlight the contribution of the assay model.

### Future directions

In future work, we plan to expand our methodology to incorporate temporal dynamics, including time-varying force of infection ([52]), maternal immunity, cross-protection or ecological interactions between serotypes ([53, 54]), and antibody boosting as a function of repeated infections ([55]). One interesting line of investigation involves considering whether *ϕ* could serve as a correlate of protection, rather than a simple marker of past infection. Our existing framework cannot on its own establish a protective relationship: doing so would require longitudinal data linking pre-exposure antibody concentration estimates to observed outcomes. The first stage of our framework would transfer directly to such a study, supplying per-individual antibody concentration estimates in place of censored endpoint titres to a correlate-of-protection analysis, allowing the uncertainty to be propagated into an estimated protection curve.

Our methods and output enrich the possibilities of serocatalytic models and are applicable beyond the specific investigation of enteroviruses we presented here. They may be particularly useful in settings with low sample volume or sparse dilution steps, where traditional titre estimation methods are unreliable. We envision these tools eventually being integrated into large-scale serological surveys and vaccine evaluation pipelines to enhance the quality of evidence-based public health policies.

## Acknowledgements

We thank the three anonymous reviewers, whose careful reading and constructive comments greatly added to the clarity and rigour of this manuscript.

## Data Availability Statement

The analysis scripts and data used in this study are available at: https://github.com/t-refae/anti-sero.

## Author contribution

Conceptualisation (TAA, MP-S, CAD, BL, EK); Data Curation (TAA, EK); Formal Analysis (TAA, BL, EK); Methodology (TAA, MP-S, CAD, BL, EK); Supervision (MP-S, CAD, BL, EK); Writing – Original Draft Preparation (TAA, EK); Writing – Review & Editing (TAA, MP-S, CAD, BL, EK)

## SUPPORTING INFORMATION

### Methods

#### Virus neutralising assays

In practice, a serum sample is diluted several times with cell-free medium, then mixed with a solution of known virus quantity and added to cell cultures in a microplate. As the virus replicates and spreads, changes in cell morphology, structure and function are observed, leading to cell death or CPE. If neutralising antibodies are present and effective in the serum dilution, then viral replication and spread will be inhibited, resulting in a reduction or elimination of CPE (see illustration in **Fig. 1**).

#### Methods for calculating 50% endpoint titre

The Reed and Muench method determines the endpoint titre as the sum of logarithms of the lower dilution (below 50% mortality) and the product of proportionate distance with the logarithms of dilution factor ([16]), as we demonstrate below. The method has been adopted for calculating the TCID_50_ for virus titration analysis [56, 57]. A similar method developed around the same time as the Reed and Muench method is the Spearman-Karber method [58, 59]. This second method yields the same result as Reed and Muench’s but does not calculate the proportionate distances and instead sums the proportion of surviving animals starting at the dilution with 100% mortality. Several improvements to these two classical methods have been proposed. A new method published in 2016 [60] does not calculate the difference of logarithms but rather, using the Reed and Muench’s mice analogy, sums the proportion of deaths across the dilutions and multiplies by the logarithm of dilution factor. A recent study improved the original Spearman-Karber method by calculating standard errors to provide a 95% confidence interval, however, this method was shown to yield wide confidence intervals [61]. Two other studies ([62, 63]) used probit or logit-log regression but do not capture the probability distribution of the outputs.

The Reed and Muench and the Spearman–Karber methods are commonly used to calculate the serum titre or 50% endpoint dilution in virus neutralising assays. Both methods produce comparable results and can be computed by hand or using widely available software e.g., Microsoft Excel, as demonstrated below. We now demonstrate these two methods using the hypothetical protective serum titration output in **Table S1**.

##### (i) Reed and Muench method

In their analyses, Reed and Muench abridged their data by disregarding the first three and last dilutions to counter the atypical or accidental occurrence of survival at high dilutions or of deaths at low dilutions. They assumed that protection from the infection challenge at a given pathogen concentration dilution step d_i_ would still have occurred at a dilution step *d*_*j*_ for all *d*_*j*_ < *d*_*i*_ [16]. That is, protection from infection at pathogen concentration dilution of 1:16 would still have occurred at the dilution of 1:8 and below. The tabulations of abridged data to calculate the endpoint titre then started at 1:8 up to 1:128, where percent mortality now starts at 0. The endpoint dilution is then calculated as follows:

log_10_ 50% endpoint dilution = log_10_ of dilution showing a mortality next below 50% + (difference of logarithms or “proportionate distance” × logarithm of dilution factor).

Difference of logarithms = [50%-(mortality at dilution next below 50%)]/[(mortality next above 50%)-(mortality next below 50%)], which is, [50 − 20]/[60 − 20] = 30/40 = 0.75

log_10_ of dilution showing a mortality next below 50% = log_10_ (16) = 1.2

log_10_ of dilution factor = log_10_ (2) = 0.3

Therefore, log_10_ 50% endpoint dilution = 1.2 + (0.75 * 0.3) = 1.425, and 50% endpoint dilution = 10 ^1.425^ = 26.9.

##### (ii) Spearman–Karber method

The endpoint dilution for this method is calculated as follows:

log_10_ 50% endpoint dilution = (*x*_0_ − *d*/2 + *d* + ∑ *r*_*i*_/*n*_*i*_)

where, *x*_0_ = log_10_ of the highest dilution at which all replicates are alive (no deaths); d = log_10_ of the dilution factor; *n*_*i*_ = number of replicates in each dilution; *r*_*i*_ = number alive (out of *n*_*i*_) in each dilution starting at *x*_0_.

Using the data in **Table S1**,

*x*_0_ = log_10_(8) = 0.9; *d* = log_10_ (2) = 0.3; *n*_*i*_ = 6; *r*_*i*_ are the values in column ‘Alive’;

Therefore, log_10_ 50% endpoint dilution = 1.45; 50% endpoint dilution = 10 ^1. 45^ = 28.5.

#### Enterovirus serum titrations used for estimation of neutralising antibody concentration

The enterovirus serology data were derived from serum samples collected as an approximately representative age-stratified cross section of the UK population. These samples had initially been selected for an EV-D68 seroepidemiology study and were collected in (i) 2006, before the reports of increased number of EV-D68 cases (n = 517), (ii) in 2011 (n = 504) and (iii) in 2017 after the 2014 EV-D68 widespread outbreaks (n = 567). The samples were obtained from the seroepidemiology unit archive collection of Public Health England (PHE, now United Kingdom Health Security Agency; Manchester, UK). The archive is an opportunistic collection of residual clinical samples from laboratories throughout England. All samples were anonymised and any patient identifying information unlinked. We only retained age and date of collection information for analyses described here.

For serum titrations, we used a standard microneutralisation assay protocol, which included inactivation of serum followed by preparation of serial 2-fold dilutions of serum in a range from 1:8 to 1:1024. The dilutions were prepared in two replicates and mixed with a solution of virus particles and that of monolayer cells. The monolayer cells were scored for CPE indicating the presence of non-neutralised virus or absence of neutralising antibodies. Serum antibody titres were recorded as the highest serum dilution preventing CPE / virus replication.

#### Choice of *k*_0_

In Eq. (8), we chose *k*_0_ = 0 such that *π*_*i*_(*d*) = Φ(0) = 0.5 when *x* = *x*_ref_, to simplify the parametrisation and improve identifiability. Suppose we instead choose a reference value 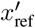 such that *k*_0_ = *ζ* ≠ 0, such that our new antibody concentration estimates for individual *i* are

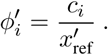

Then, for all individuals *i*, we have

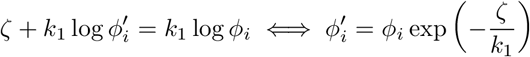

Since this rescaling is common to all individuals and does not depend on the dilution, the linear predictor is unchanged,

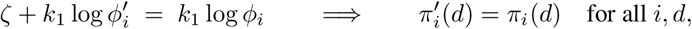

and hence the two parametrisations index the same likelihood,

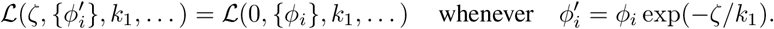

We confirm this empirically in **Fig. S1** by refitting the first-stage assay model with *k*_0_ = ±1, ±2.

#### Detailed analytical solutions for the serocatalytic models

Eq. (12a) is a separable first-order ODE whose solution for a given age *a* is:

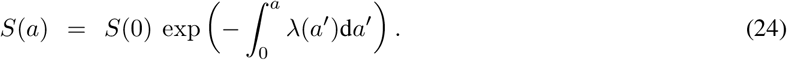

Eq. (12b) can be solved with an integrating factor facilitator function:

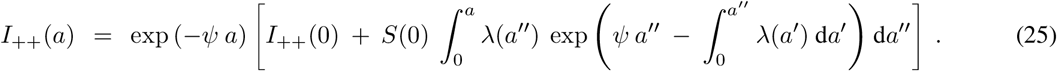

The analytical steps taken to derive Eq. (25) are outlined below. Using the integrating factor facilitator function approach on Eq. (12b) yields:

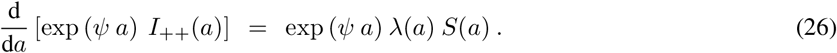

Integrating both sides, we get:

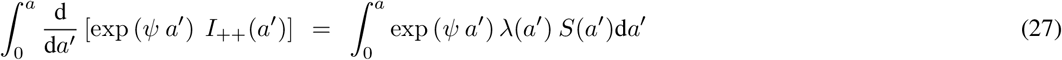

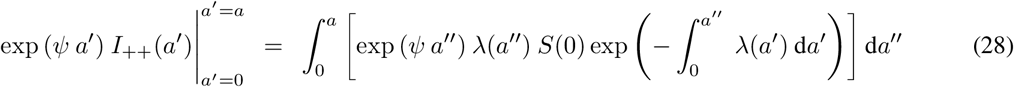

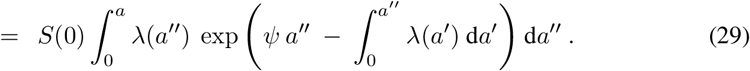

Evaluating the left-hand side and rearranging, we arrive at the full analytical solution for the *I*_++_ compartment given in Eq. (25).

The analogous equation in the case where the force of infection 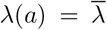 is assumed fixed for some age *a* ∈ [*t* − *τ, t* − *τ* + 1] is given by:

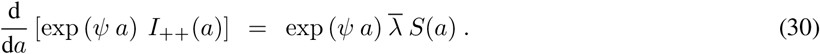

Integrating between *a* = *t* − *τ* and *a* = *t* − *τ* + 1 yields:

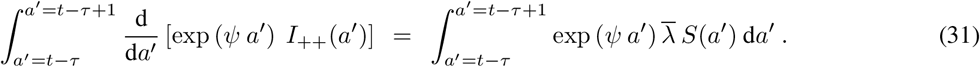

If we assume that the force of infection *λ*(*a*) is piecewise constant between 0 and *a* and note that for some *a* ∈ [*t* − *τ, t* − *τ* + 1]:

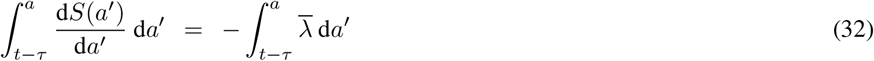

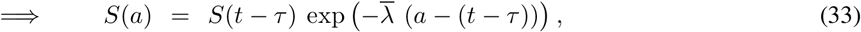

then:

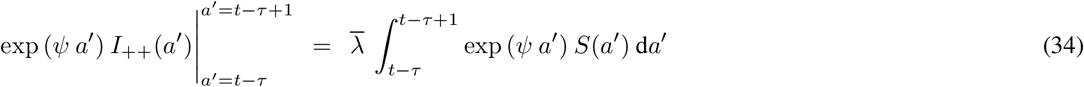

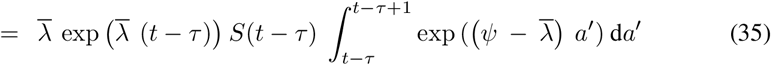

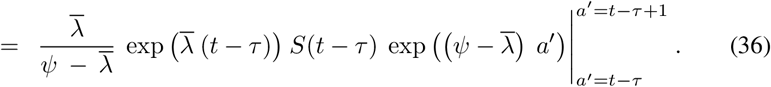

Rearranging for *I*_++_(*t* − *τ* + 1) on the left-hand side and substituting *a* := *t* − *τ* we get:

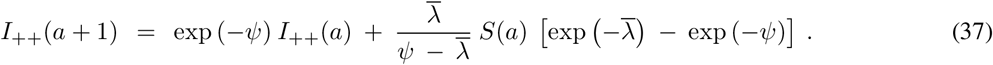

When 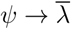, the limit exists and is equal to:

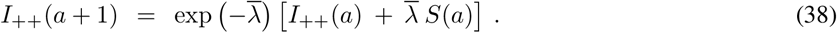

## Supporting Tables

**Table S1:** Reed and Muench’s 1938 “Titration of a hypothetical protective serum” data. The Reed and Muench method also addresses the occurrence of ‘accidental’ survivals and deaths, where presence or absence of cytopathic effects in cell monolayers are occasionally encountered across a series of dilutions (for example, see dilutions 8 and 256). The values in the ‘Cumulative survival’ column are sums of ‘Alive’ column starting from the bottom-to-top. The values in the ‘Cumulative deaths’ column are sums of ‘Dead’ column starting from the top-to-bottom.

| Dilution | Alive | Dead | Survival | Cumulative survival | Cumulative deaths | Percent mortality |
| --- | --- | --- | --- | --- | --- | --- |
| 1:1 | 6 | 0 | 100% | 32 | 0 | 0 |
| 1:2 | 6 | 0 | 100% | 26 | 0 | 0 |
| 1:4 | 5 | 1 | 83% | 20 | 1 | 5 |
| 1:8 | 6 | 0 | 100% | 15 | 1 | 6 |
| 1:16 | 4 | 2 | 67% | 9 | 3 | 25 |
| 1:32 | 2 | 4 | 33% | 5 | 7 | 58 |
| 1:64 | 2 | 4 | 33% | 3 | 11 | 79 |
| 1:128 | 0 | 6 | 0% | 1 | 17 | 94 |
| 1:256 | 1 | 5 | 17% | 1 | 22 | 96 |

**Table S2:** Convergence summary table across enteroviruses, model types, and *σ*_*ϕ*_ specifications. A checkmark ✓ (cross ×) denotes a model fit that did (not) converge. Convergence was assessed against the criteria detailed in **Section 1.8**.

| Enterovirus | States | Seroreversion? | Shared $\sigma_\phi$ | Serostate-specific $\sigma_\phi$ | Total converged |
| --- | --- | --- | --- | --- | --- |
| CVA6 | 2 | No (SI) | ✓ | ✓ | 3 |
|  |  | Yes (SIS) | × | × |  |
|  | 3 | No (SII) | ✓ | × |  |
|  |  | Yes (SIIS) | × | × |  |
| EV-A71 | 2 | No (SI) | ✓ | ✓ | 5 |
|  |  | Yes (SIS) | × | × |  |
|  | 3 | No (SII) | ✓ | ✓ |  |
|  |  | Yes (SIIS) | ✓ | × |  |
| EV-D68 | 2 | No (SI) | ✓ | ✓ | 4 |
|  |  | Yes (SIS) | ✓ | × |  |
|  | 3 | No (SII) | ✓ | × |  |
|  |  | Yes (SIIS) | × | × |  |
| Total converged |  |  | 8 | 4 | 12 |

**Table S3:**
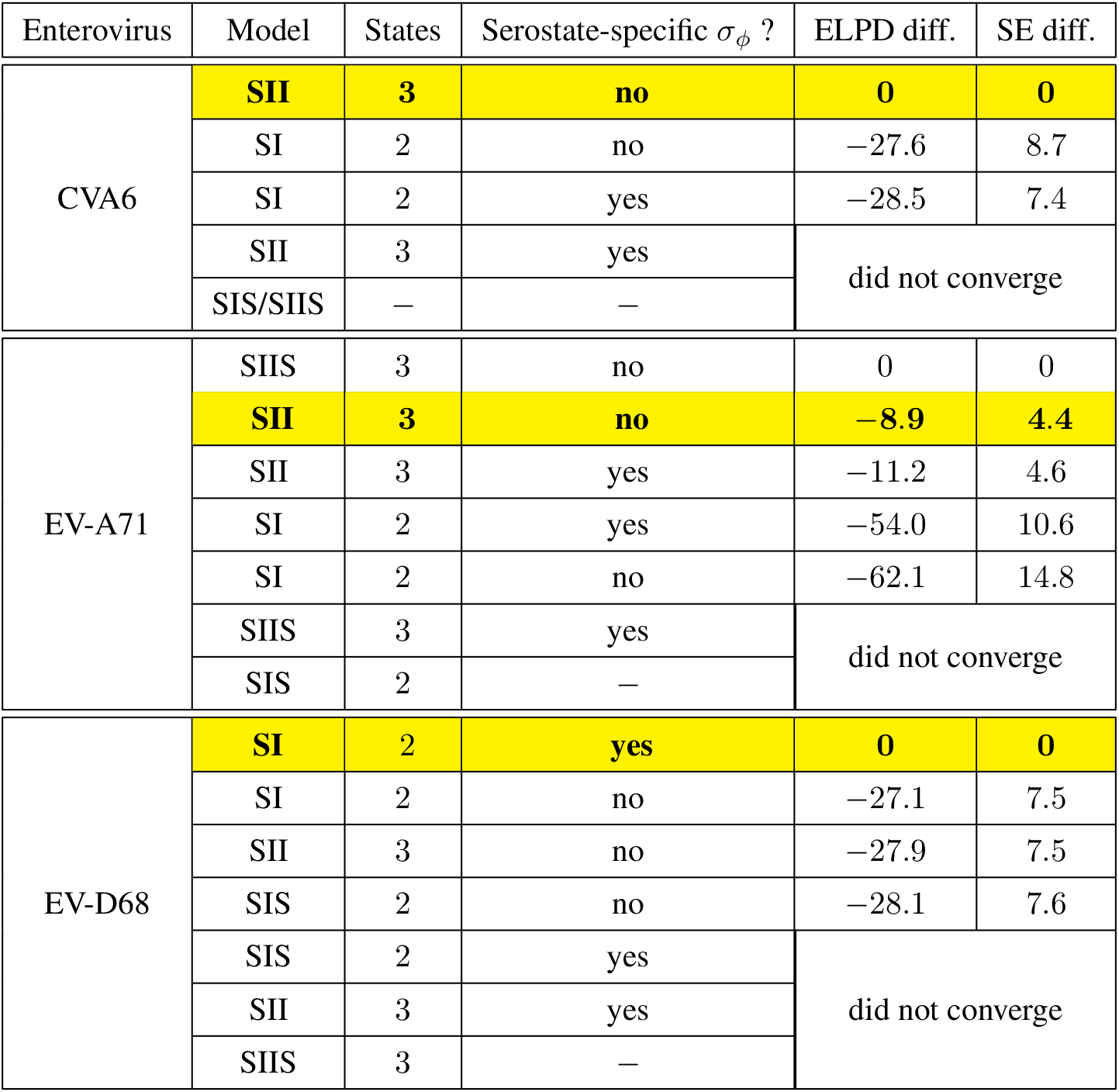
Approximate LOO-CV comparison of model structures (SI/SII/SIS/SIIS, each with shared or serostate-specific *σ*_*ϕ*_) for each enterovirus serotype. Expected log predictive density differences (ELPD diff.) are reported relative to the best-scoring converged model for that serotype, with the associated standard error (SE diff.), computed with approximate leave-one-out cross-validation via the R package loo [32]. Only fits meeting the convergence criteria in **Section 1.8** were compared. Highlighted rows denote the specification adopted as our primary analysis. For CVA6 and EV-D68 this is the best-scoring model. For EV-A71, we retain the SII model as our primary specification for comparability with CVA6 (where *ω* was not identifiable) and report the SIIS results as a sensitivity analysis (see **Results**).

| Enterovirus | Model | States | Serostate-specific $\sigma_\phi$ ? | ELPD diff. | SE diff. |
| --- | --- | --- | --- | --- | --- |
| CVA6 | **SII** | **3** | **no** | **0** | **0** |
|  | SI | 2 | no | -27.6 | 8.7 |
|  | SI | 2 | yes | -28.5 | 7.4 |
|  | SII | 3 | yes | did not converge |  |
|  | SIS/SIIS | — | — |  |  |
| EV-A71 | SIIS | 3 | no | 0 | 0 |
|  | **SII** | **3** | **no** | **-8.9** | **4.4** |
|  | SII | 3 | yes | -11.2 | 4.6 |
|  | SI | 2 | yes | -54.0 | 10.6 |
|  | SI | 2 | no | -62.1 | 14.8 |
|  | SIIS | 3 | yes | did not converge |  |
|  | SIS | 2 | — |  |  |
| EV-D68 | **SI** | **2** | **yes** | **0** | **0** |
|  | SI | 2 | no | -27.1 | 7.5 |
|  | SII | 3 | no | -27.9 | 7.5 |
|  | SIS | 2 | no | -28.1 | 7.6 |
|  | SIS | 2 | yes | did not converge |  |
|  | SII | 3 | yes |  |  |
|  | SIIS | 3 | — |  |  |

**Table S4:** Parameter estimates for EV-A71 under the SII and SIIS specifications. Posterior medians and central 95% credible intervals for the serocatalytic model parameters, estimated under our primary specification for EV-A71 (SII, shared *σ*_*ϕ*_) and under the corresponding model including an *I*_+_ → *S* transition at rate *ω* (SIIS, shared *σ*_*ϕ*_), the only model incorporating seroreversion to converge for this serotype. *µ*_S_, *µ*_*I*++_ and *µ*_*I*+_ are the location parameters of the lognormal antibody concentration distribution in each serostate, *σ*_*ϕ*_ the scale parameter shared across serostates, *ψ* the serological waning rate, *ω* the seroreversion rate, and *λ*_1_ and *κ* the parameters of the age-dependent force of infection in Eq. (23). The seroreversion half-life *t*_1/2_ = log 2/*ω* is derived from the posterior draws of *ω* and applies to the SIIS model only. 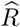 is the rank-normalised statistic and ESS the bulk effective sample size [31]; both fits met our convergence criteria of 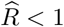 and ESS > 400 on all parameters. Allowing seroreversion lowers the estimated rate of decline of the force of infection with age and raises the inferred susceptible proportion in adulthood; the corresponding serodynamics are shown in **Fig. S7**.

| EV-A71: SII and SIIS parameter estimates |  |  |  |  |  |  |  |  |
| --- | --- | --- | --- | --- | --- | --- | --- | --- |
|  | SII (primary) |  |  |  | SIIS (sensitivity) |  |  |  |
|  | MEDIAN | 95% CrI | R-HAT | ESS | MEDIAN | 95% CrI | R-HAT | ESS |
| $\mu_S$ | 2.71 | 2.65 – 2.77 | 1.002 | 2950 | 2.72 | 2.66 – 2.78 | 1.000 | 2586 |
| $\mu_{I_{++}}$ | 6.34 | 6.20 – 6.50 | 1.001 | 2381 | 6.38 | 6.22 – 6.54 | 1.002 | 3889 |
| $\mu_{I_+}$ | 4.73 | 4.49 – 4.96 | 1.002 | 1242 | 4.81 | 4.58 – 5.05 | 1.001 | 1659 |
| $\sigma_\phi$ | 0.763 | 0.722 – 0.808 | 1.001 | 2679 | 0.763 | 0.723 – 0.807 | 1.000 | 2641 |
| $\psi$ | 0.038 | 0.028 – 0.050 | 1.001 | 1679 | 0.049 | 0.036 – 0.070 | 1.002 | 2270 |
| $\omega$ | – | – | – | – | 0.013 | 0.0071 – 0.020 | 1.000 | 2452 |
| $\lambda_1$ | 0.157 | 0.116 – 0.218 | 1.001 | 2045 | 0.133 | 0.099 – 0.183 | 1.001 | 1988 |
| $\kappa$ | 0.342 | 0.230 – 0.551 | 1.001 | 1855 | 0.208 | 0.131 – 0.338 | 1.002 | 1824 |
| $t_{1/2}$ (years) | – | – | – | – | 54.0 | 34.8 – 97.6 | 1.000 | 2452 |

**Table S5:** Summary statistics of fitted parameters for three enterovirus serotypes. With respect to the posterior distributions of the variables for each serotype: “SD” denotes the standard deviation; “Q2.5” and “Q97.5” are the lower and upper bounds of the central 95% credible interval, respectively; “RHAT” denotes the rank-normalised 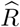 metric used to evaluate MCMC convergence; “ESS_BULK” (“ESS_TAIL”) is the bulk (tail) effective sample size, a diagnostic to assess the sampling efficiency in the bulk (tail) of the posterior distribution [31].

**CVA6: Summary Statistics of Fitted Parameters**
| VARIABLE | MEAN | MEDIAN | SD | Q2.5 | Q97.5 | RHAT | ESS_BULK | ESS_TAIL |
| --- | --- | --- | --- | --- | --- | --- | --- | --- |
| $\mu_S$ | 2.948 | 2.947 | 0.048 | 2.854 | 3.045 | 1.002 | 2,423 | 1,883 |
| $\mu_{I_{++}}$ | 6.147 | 6.145 | 0.111 | 5.938 | 6.365 | 1.000 | 2,641 | 2,344 |
| $\mu_{I_{+}}$ | 4.868 | 4.877 | 0.182 | 4.489 | 5.193 | 1.001 | 1,396 | 1,364 |
| $\sigma_\phi$ | 0.922 | 0.921 | 0.029 | 0.869 | 0.981 | 1.002 | 1,680 | 1,953 |
| $\psi$ | 0.045 | 0.043 | 0.013 | 0.024 | 0.078 | 1.001 | 1,958 | 2,114 |
| $\lambda_1$ | 0.266 | 0.263 | 0.042 | 0.195 | 0.356 | 1.006 | 1,734 | 1,804 |
| $\kappa$ | 0.486 | 0.469 | 0.120 | 0.307 | 0.760 | 1.005 | 1,742 | 1,700 |

**EV-A71: Summary Statistics of Fitted Parameters**

| VARIABLE | MEAN | MEDIAN | SD | Q2.5 | Q97.5 | RHAT | ESS_BULK | ESS_TAIL |
| --- | --- | --- | --- | --- | --- | --- | --- | --- |
| $\mu_S$ | 2.708 | 2.708 | 0.032 | 2.646 | 2.769 | 1.002 | 2,950 | 2,417 |
| $\mu_{I_{++}}$ | 6.345 | 6.343 | 0.076 | 6.200 | 6.497 | 1.001 | 2,381 | 2,277 |
| $\mu_{I_{+}}$ | 4.730 | 4.733 | 0.118 | 4.491 | 4.961 | 1.002 | 1,242 | 1,527 |
| $\sigma_\phi$ | 0.763 | 0.763 | 0.022 | 0.722 | 0.808 | 1.001 | 2,679 | 2,129 |
| $\psi$ | 0.038 | 0.038 | 0.005 | 0.028 | 0.050 | 1.001 | 1,679 | 1,884 |
| $\lambda_1$ | 0.159 | 0.157 | 0.027 | 0.116 | 0.218 | 1.001 | 2,045 | 1,514 |
| $\kappa$ | 0.355 | 0.342 | 0.084 | 0.230 | 0.551 | 1.001 | 1,855 | 1,362 |

**EV-D68: Summary Statistics of Fitted Parameters**

**Table S5: Summary statistics of fitted parameters for three enterovirus serotypes.**
| VARIABLE | MEAN | MEDIAN | SD | Q2.5 | Q97.5 | RHAT | ESS_BULK | ESS_TAIL |
| --- | --- | --- | --- | --- | --- | --- | --- | --- |
| $\mu_S$ | 4.346 | 4.348 | 0.070 | 4.208 | 4.480 | 1.001 | 1,587 | 1,907 |
| $\mu_{I_{++}}$ | 6.205 | 6.207 | 0.065 | 6.068 | 6.328 | 1.002 | 1,657 | 1,893 |
| $\sigma_{\phi,S}$ | 1.272 | 1.271 | 0.043 | 1.189 | 1.362 | 1.001 | 1,332 | 1,961 |
| $\sigma_{\phi,I_{++}}$ | 0.833 | 0.830 | 0.051 | 0.739 | 0.938 | 1.004 | 1,413 | 1,794 |
| $\lambda_1$ | 0.060 | 0.059 | 0.008 | 0.046 | 0.077 | 1.003 | 1,590 | 1,901 |
| $\kappa$ | 0.017 | 0.018 | 0.008 | 0.002 | 0.033 | 1.003 | 1,299 | 1,094 |

## Supporting Figures

**Figure S1:**
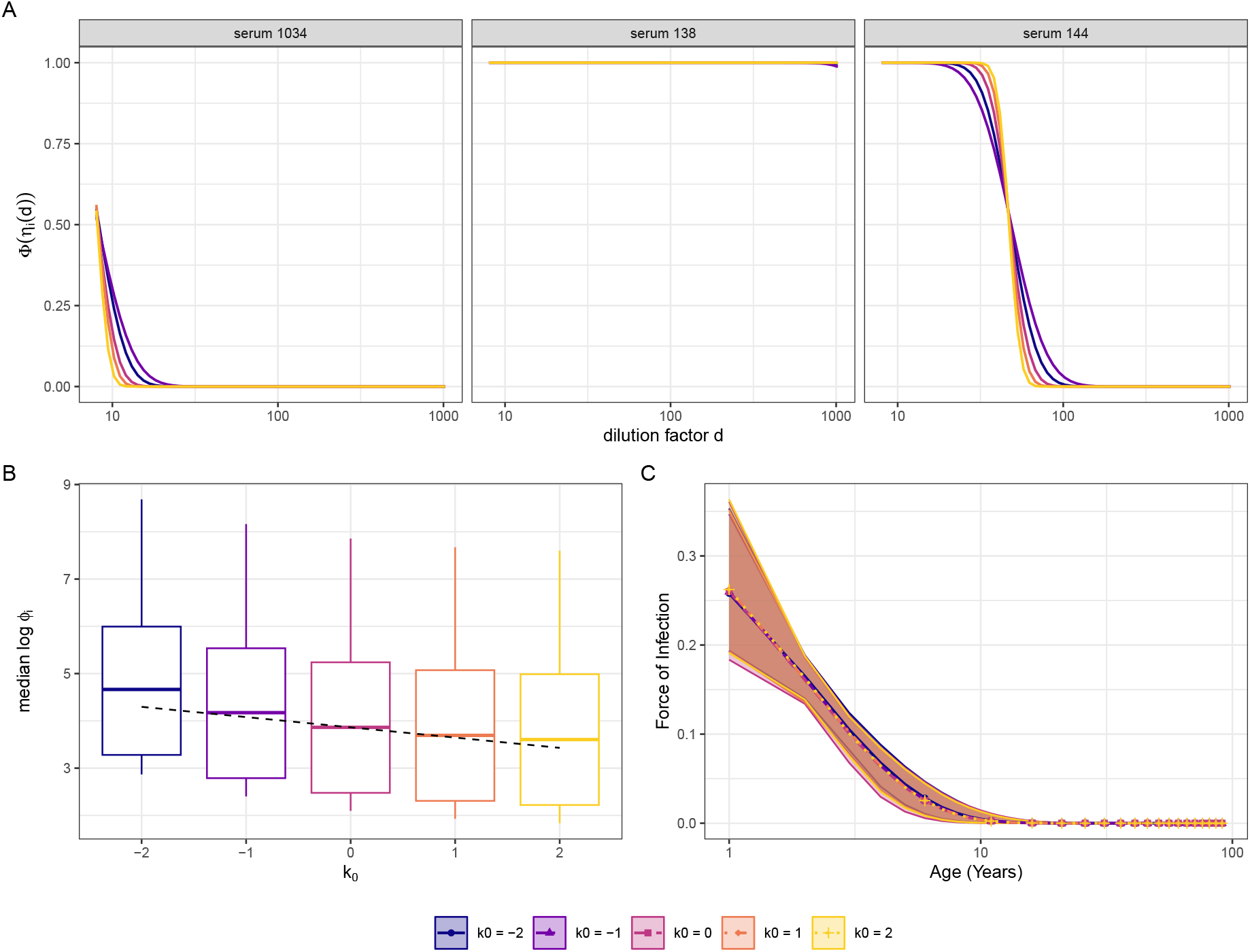
Invariance of the first-stage model to the choice of *k*_0_, illustrated for CVA6. The first-stage assay model was refitted with *k*_0_ = −2, −1, 0, 1, 2, corresponding to different choices of the reference concentration *x*_ref_. The fitted dose-response curves and the downstream force-of-infection estimates are unaffected.

**Figure S2:**
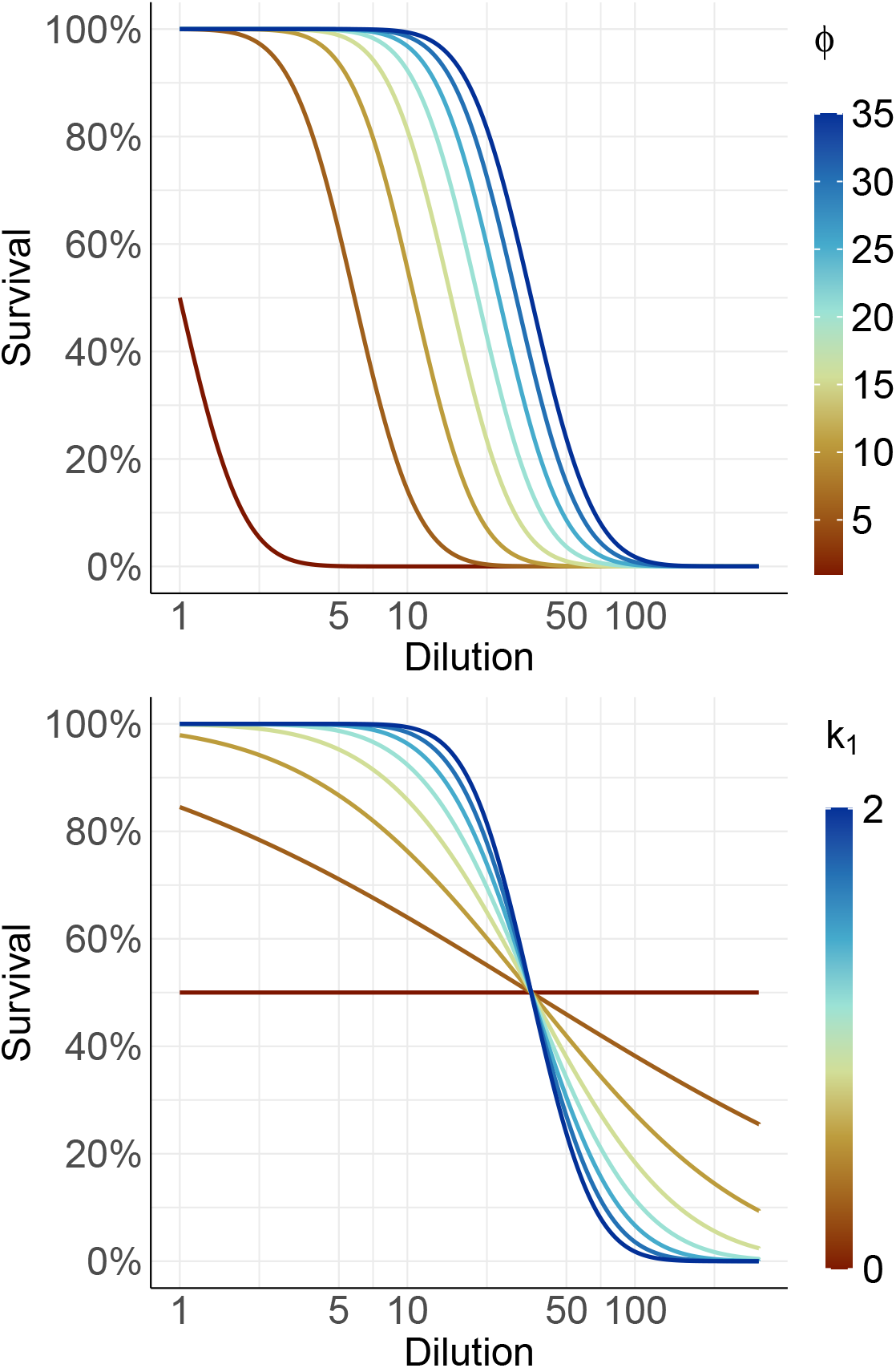
Visualising the modelled dose-response relationship. This graph shows Eq. (10) as a function of serum dilution when varying *ϕ* (top) and *k*_1_ (bottom). The colour bars on the right indicate the range of values visualised in each panel. In the top (bottom) plot, the baseline value of *k*_1_ = 2 (*ϕ* = 35) was used for visualisation purposes.

**Figure S3:**
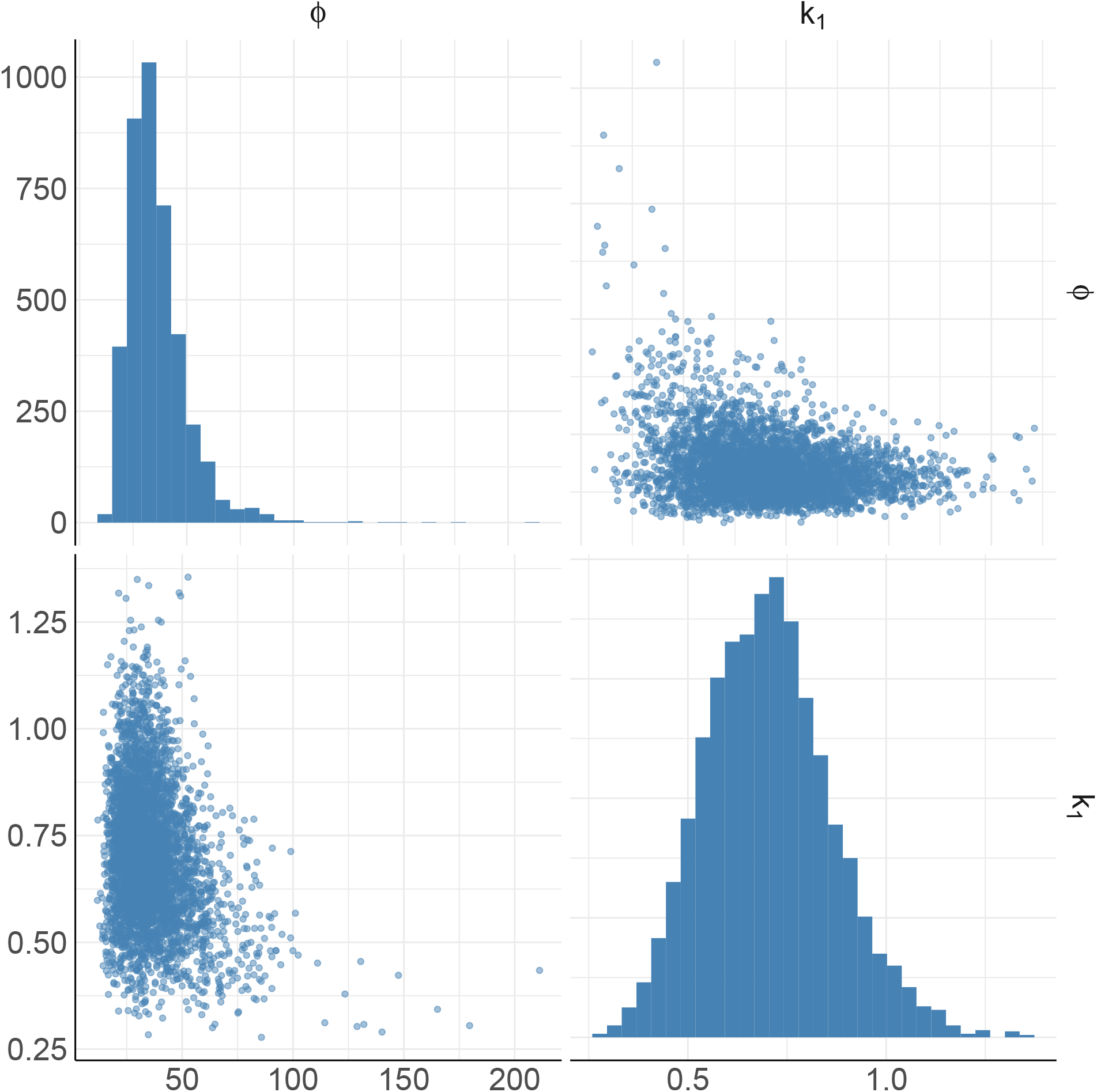
Uncertainty in the modelled estimates from fitting the Reed and Muench data Table S1. The plots on the diagonal show the marginal distributions of the two parameters in Eq. (10) (*ϕ* and *k*_1_); the off-diagonal graphs visualise the joint bivariate distributions obtained from the posterior draws.

**Figure S4:**
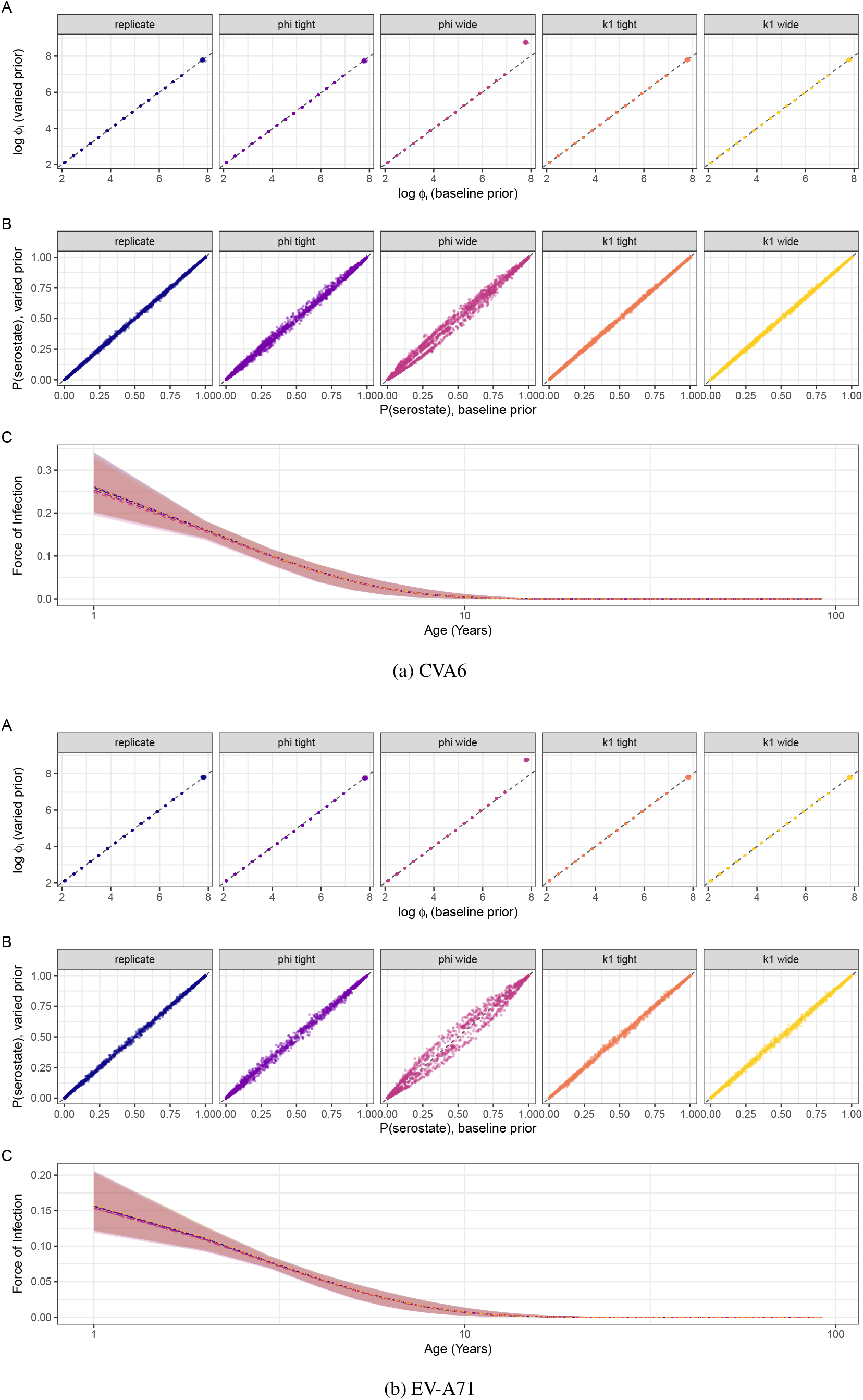
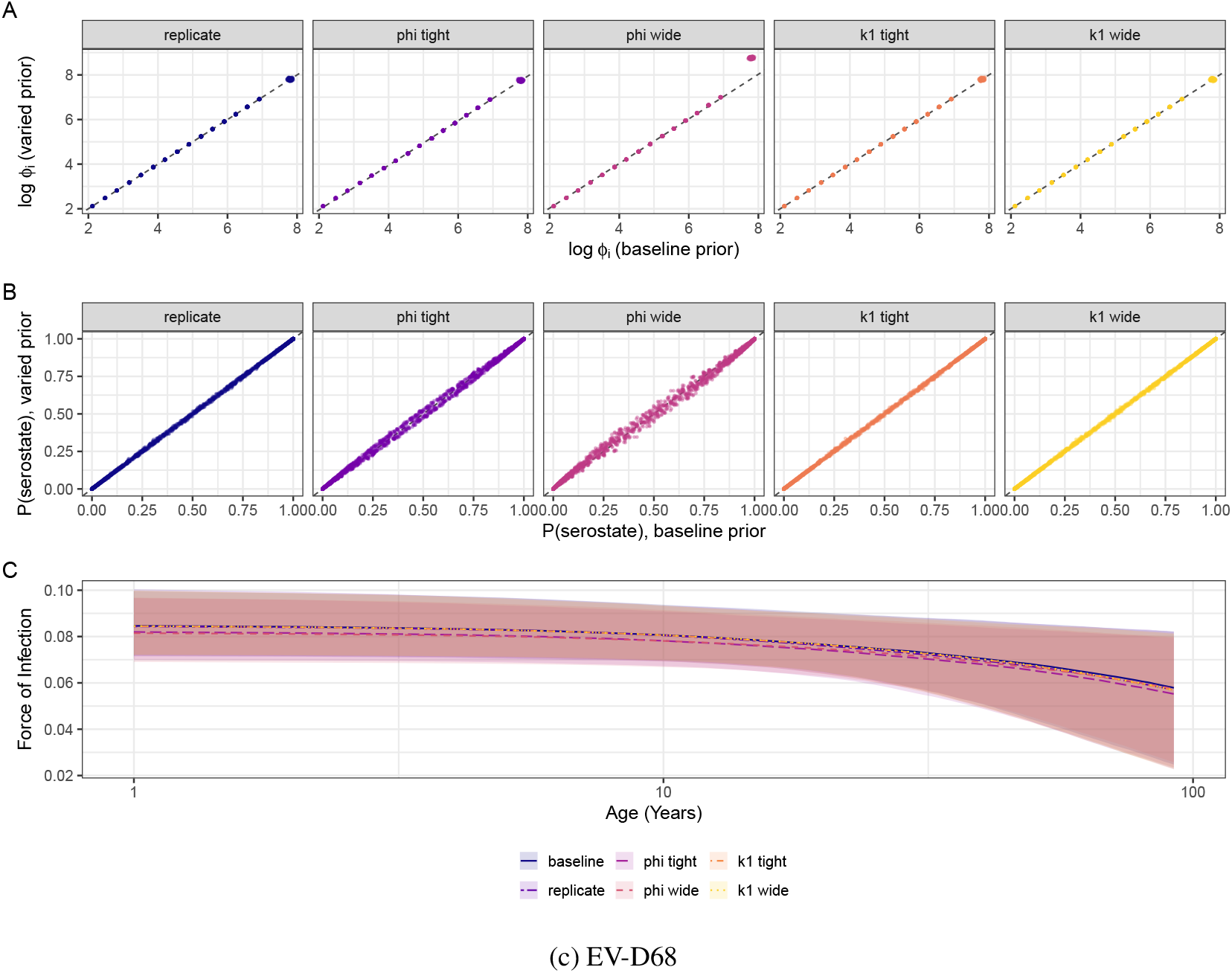
Prior sensitivity analysis for *k*_1_ and *ϕ* for CVA6, EV-A71, and EV-D68. In each panel, we compare the posterior log *ϕ*_*i*_ values (top row) and the maximum probability of serostate assignment for each individual (middle row). From left to right, the comparison is between our baseline prior distributions (given in **Table 1**) and a replicate run (same baseline prior distributions for both *ϕ* and *k*_1_), a “tight” *ϕ*-prior distribution (*ϕ* ∼ Cauchy^+^(0, 50) unchanged k_1_-prior distribution), a “wide” *ϕ*-prior distribution (*ϕ* ∼ Cauchy^+^(0, 5000); unchanged k_1_-prior distribution), a “tight” k_1_-prior distribution (*k*_1_ ∼ Cauchy^+^(0, 0.25) unchanged *ϕ*-prior distribution), and a “wide” *k*_1_-prior distribution (*k*_1_ ∼ Cauchy^+^(0, 4); unchanged *ϕ*-prior distribution). The bottom row overlays the estimated force of infection across age along with the central 95% credible interval ribbons.

**Figure S5:**
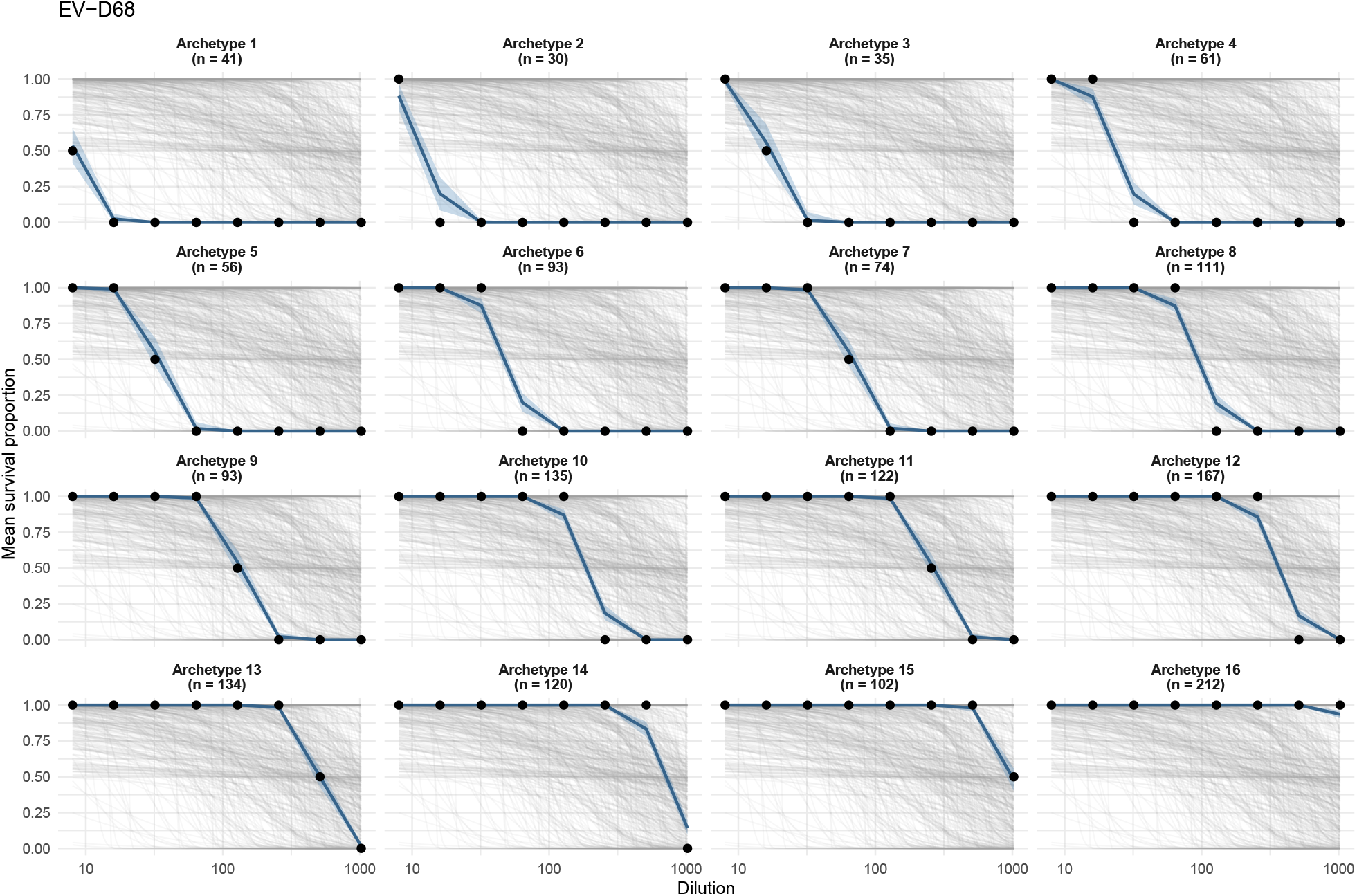
Prior and posterior predictive checks for the EV-D68 antibody concentration model. Observed EV-D68 serum titration survival outcomes are grouped by unique survival-response archetype across the dilution series. Black points show the observed mean proportion of surviving cells at each serum dilution within each archetype (two replicates per serosurveyed individual). Grey lines show prior predictive survival trajectories generated from the mechanistic antibody model prior distributions, while the blue lines and shaded ribbons convey the posterior predictive median and central 95% posterior predictive interval, respectively. The prior predictive trajectories indicate that our antibody concentration model prior distributions have the freedom to reproduce a wide range of potential survival-dilution relationships; the posterior predictive distributions broadly reproduce the observed dilution-response patterns, indicating that the probit antibody concentration model captures the observed EV-D68 titration data.

**Figure S6:**
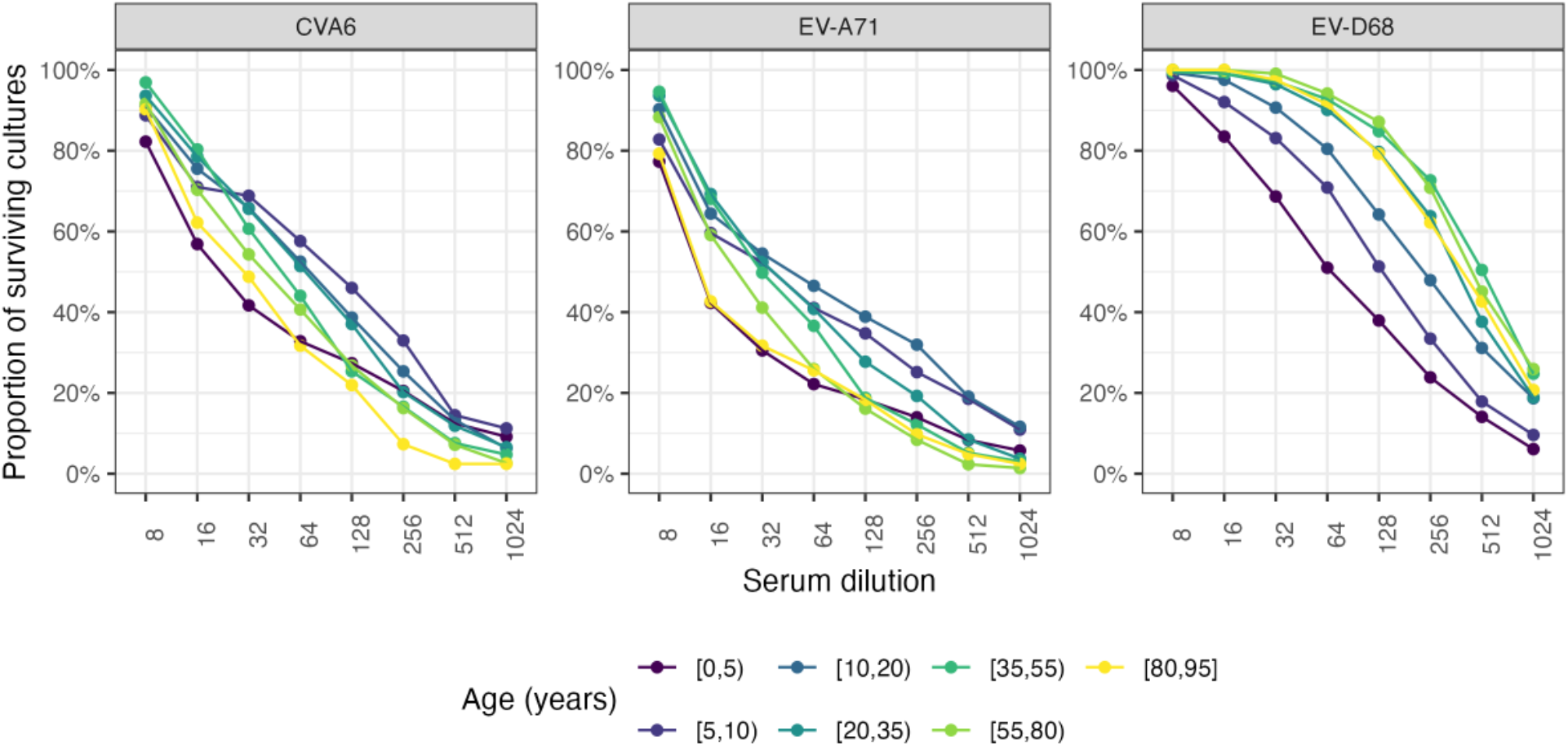
Output of serum titration against three enterovirus serotypes. This plot shows the observed proportion of surviving cell cultures (y-axis) as a function of serum dilution levels (x-axis) in different age categories. Serum samples were diluted eight times in a microplate, in 2-fold, and the presence of antibodies determined by counting the proportion of wells with surviving cell cultures per dilution level.

**Figure S7:**
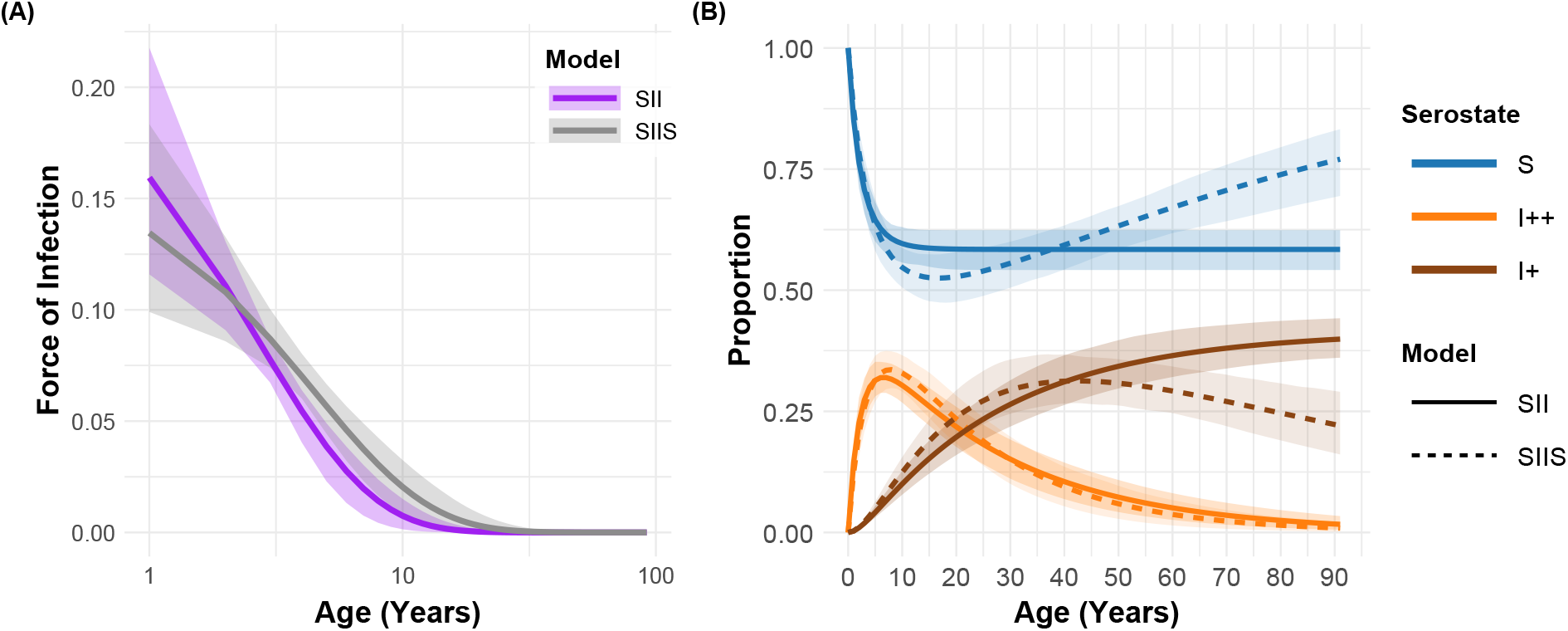
Sensitivity of the EV-A71 results to the inclusion of seroreversion. Comparison of our primary specification (SII, shared *σ*_*ϕ*_) with the corresponding model including an *I*_+_ → *S* transition at rate *ω* (SIIS, shared *σ*_*ϕ*_), the only model incorporating seroreversion to converge for this serotype. **(A)** Inferred age-dependent force of infection *λ*(*a*) under each model, with shaded areas denoting the central 95% credible intervals; the horizontal axis is on a log_10_ scale. **(B)** Inferred population-level proportions in each serostate under each model (solid lines: SII; dashed lines: SIIS), with serostates coloured as in **Fig. 5.** Shaded areas denote the central 95% credible intervals, drawn more darkly for the SII model.

**Figure S8:**
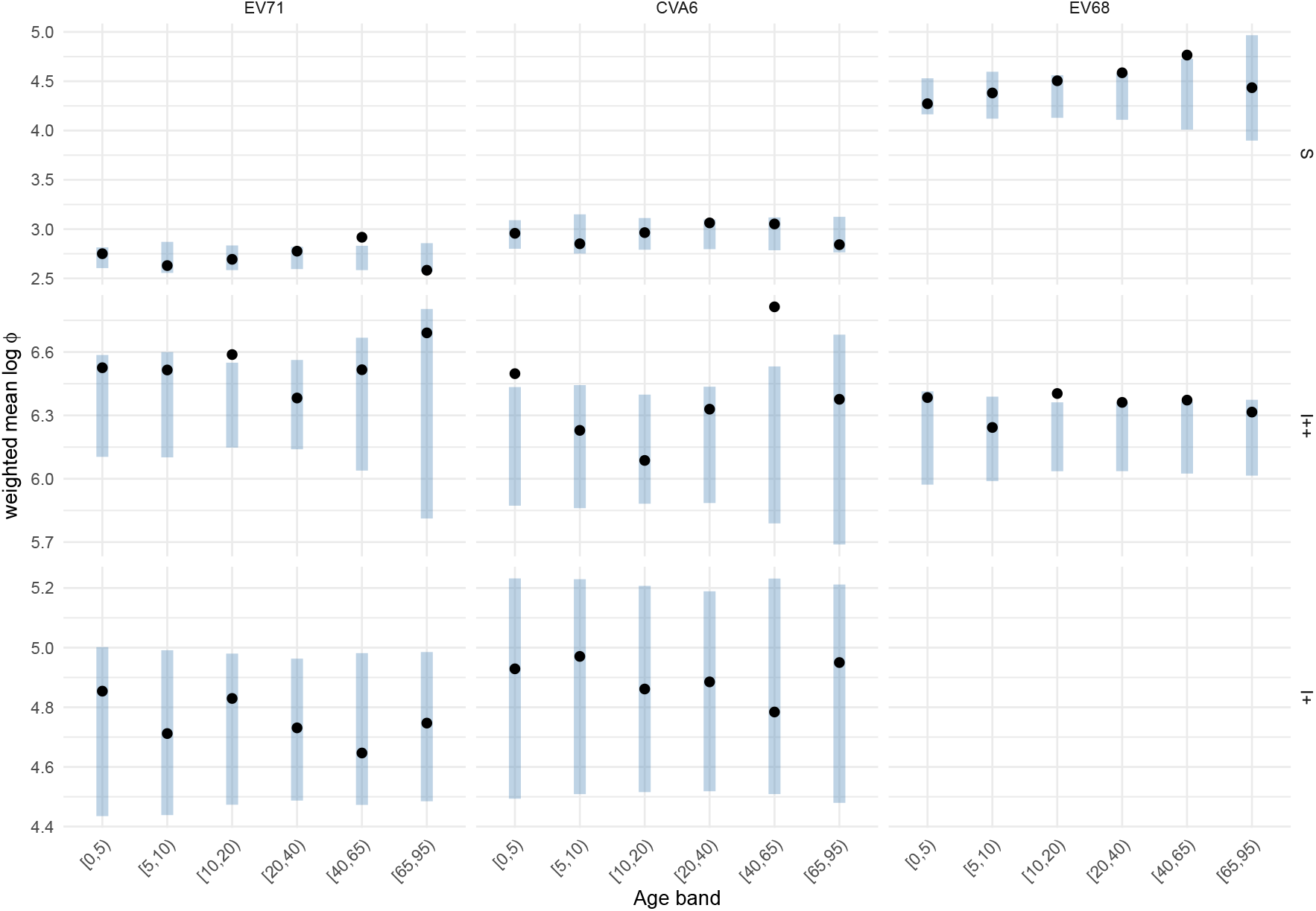
Within-serostate age profile of antibody concentration for the three enterovirus serotypes. Points show the posterior median of the serostate-weighted mean log *ϕ* in each age band, where each individual’s log *ϕ* is weighted by their posterior probability of occupying that serostate. Intervals show the central 95% posterior predictive distribution of the same quantity under the primary specification for each serotype.

**Figure S9:**
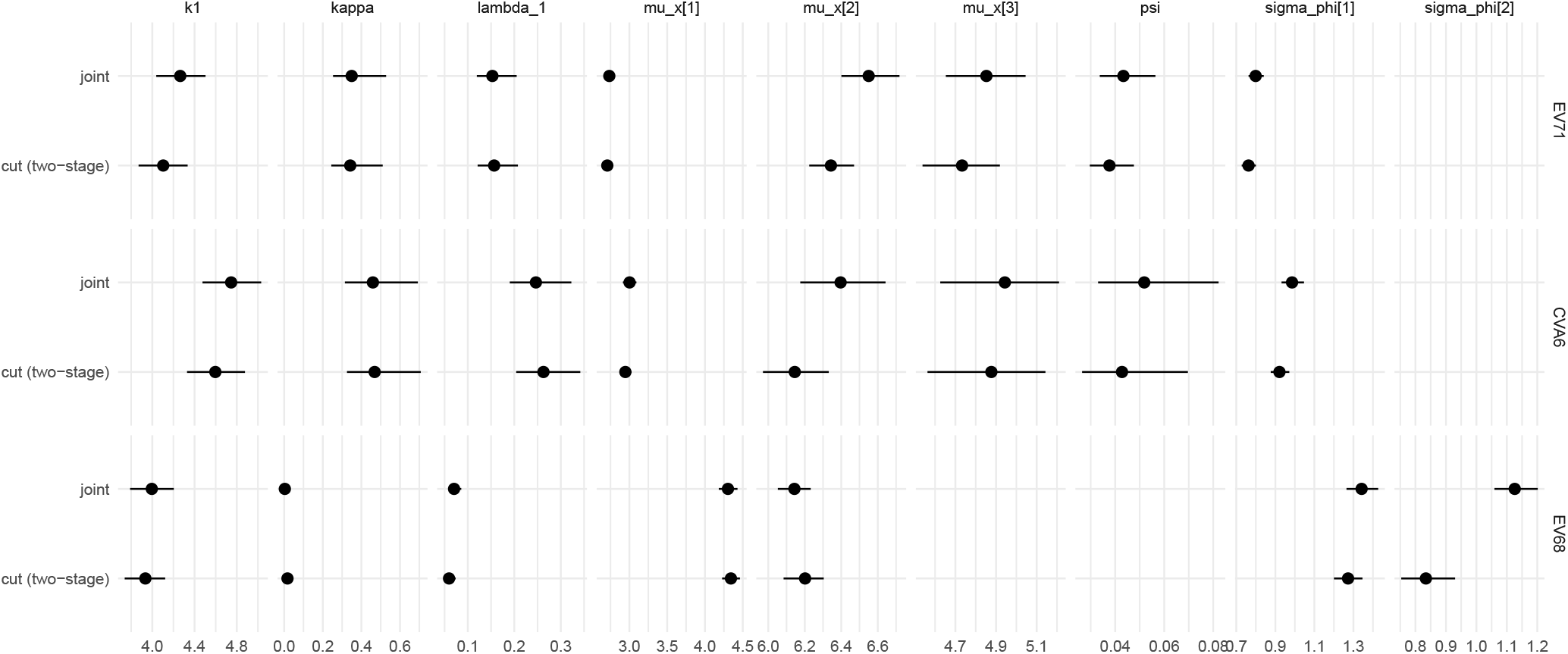
Comparison of cut and joint inference for each enterovirus serotype. Posterior medians (points) and central 95% credible intervals (bars) for each serocatalytic model parameter, estimated under our two-stage cut procedure (in which posterior draws of log *ϕ*_*i*_ are propagated from the assay model) and under a fully joint model. The comparison uses the primary specification for each serotype: SII with a shared *σ*_*ϕ*_ for CVA6 and EV-A71, and SI with serostate-specific *σ*_*ϕ*_ for EV-D68. Estimates agree closely throughout, with the sole exception of *σ*_*ϕ*,*I*++_ for EV-D68.

**Figure S10:**
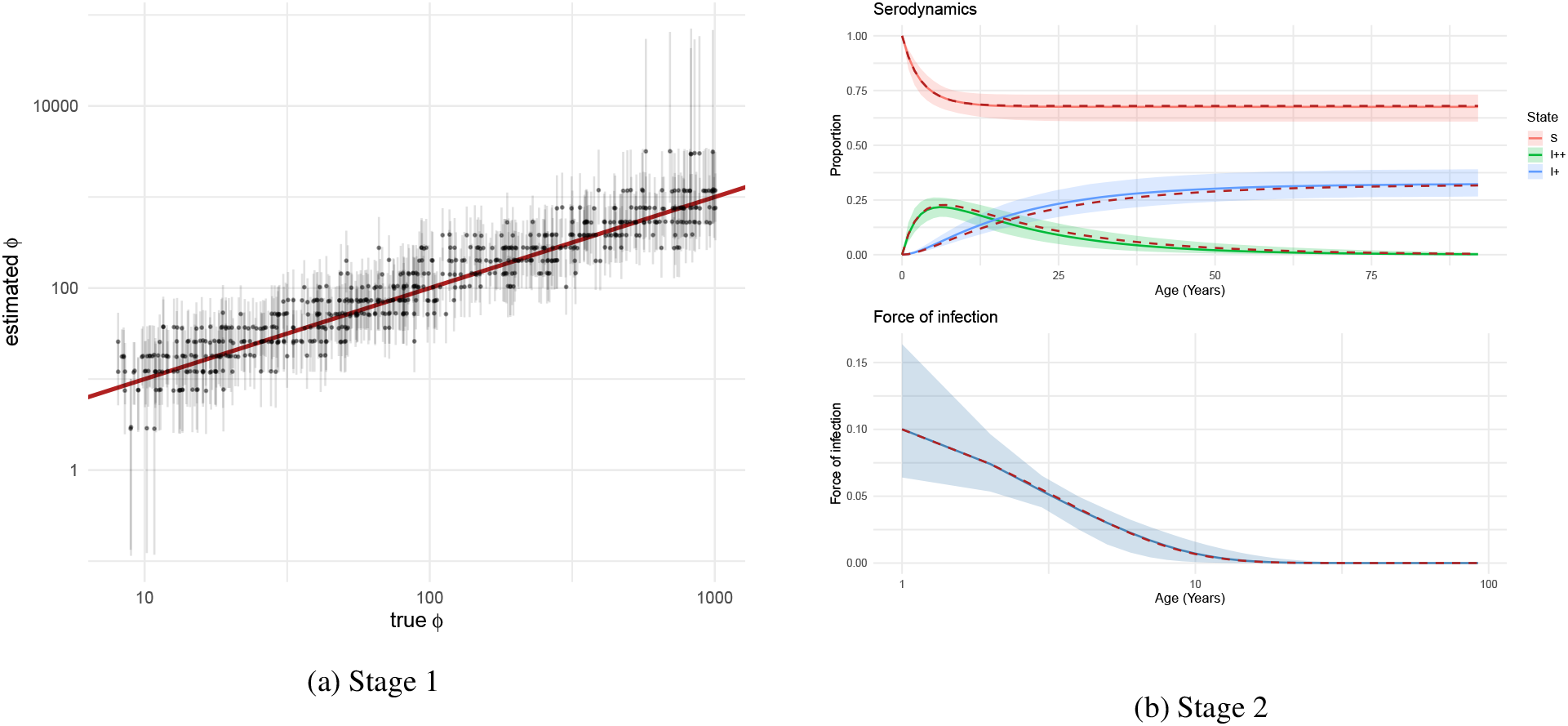
Simulation recovery study using the SII model. On the left is the log − log plot between true and estimated *ϕ* values for each individual in the first-stage fitting, with a superimposed red identity line; on the right is the resulting second-stage fitting of the serodynamics (top) and force of infection (bottom), with the simulated truth denoted by a dashed red line in all cases. The ribbon plots on the right denote the central 95% posterior credible intervals.

**Figure S11:**
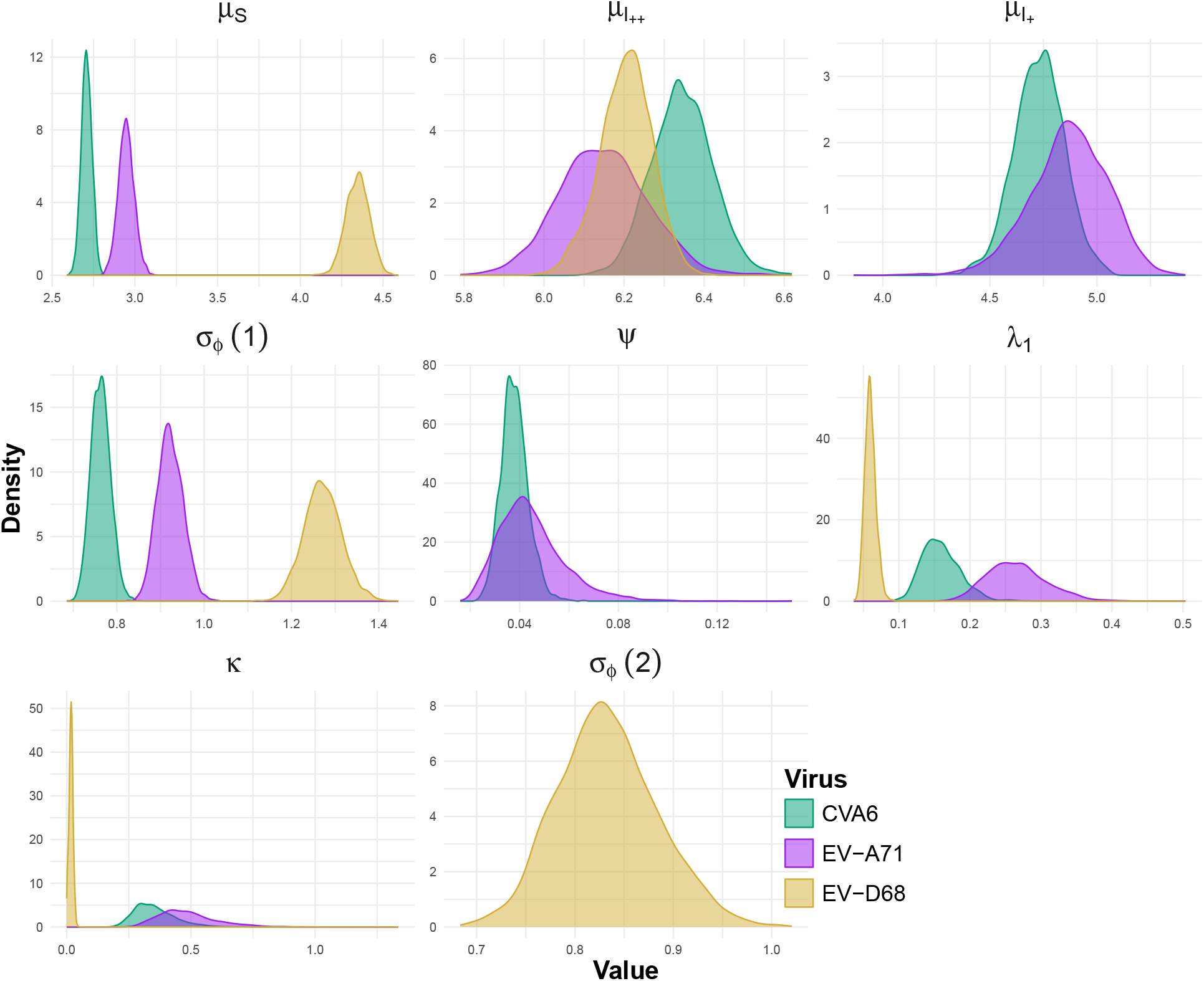
Estimated parameter values. Posterior distributions of the parameters specifying the serocatalytic model (12) for each serotype, under the primary specification in each case. The top row shows the location parameters modulating the lognormal antibody concentration distribution in each serostate: *S* –susceptible, *I*_++_ – recently infected, and *I*_+_ – infected a while ago. For EV-D68 we use the two-state SI model, which has no *I*_+_ serostate and therefore no *µ*_*I*+_ or waning rate *ψ* (see **Methods**). From left to right, the middle row shows the posterior distributions of the scale parameter *σ*_*ϕ*,*x*_ (shared across serostates for CVA6 and EV-A71; estimated separately for each serostate for EV-D68, shown as *σ*_*ϕ*_ (1) and *σ*_*ϕ*_ (2)) associated with the lognormal antibody concentration distributions, as well as the serological waning rate (*ψ*) and the force of infection parameter *λ*_1_. The bottom-left panel shows the posterior distribution of the force of infection parameter *κ*.

**Figure S12:**
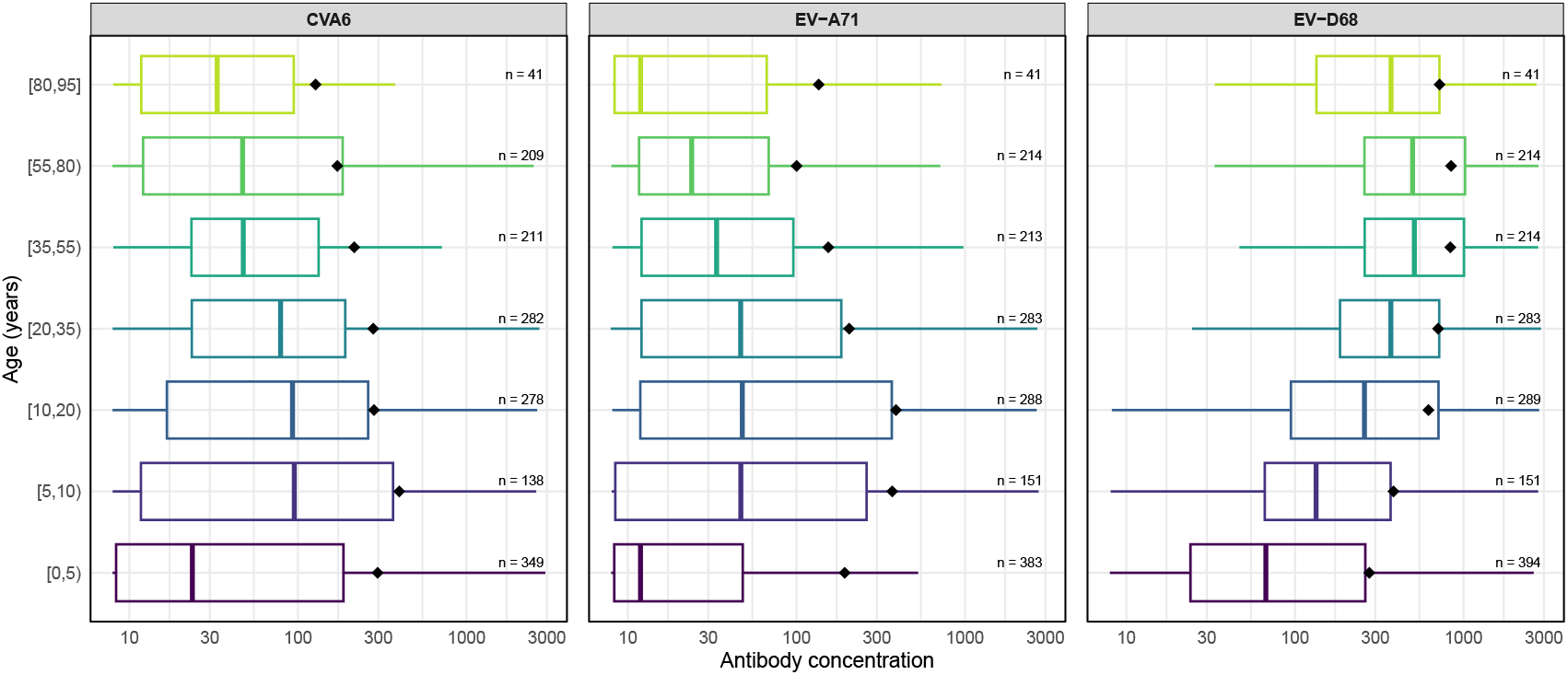
Antibody concentration against three enterovirus serotypes. This plot shows the median (coloured middle bar) and mean (black point) estimated antibody concentration (*ϕ*, x-axis) as a function of age (y-axis).

**Figure S13:**
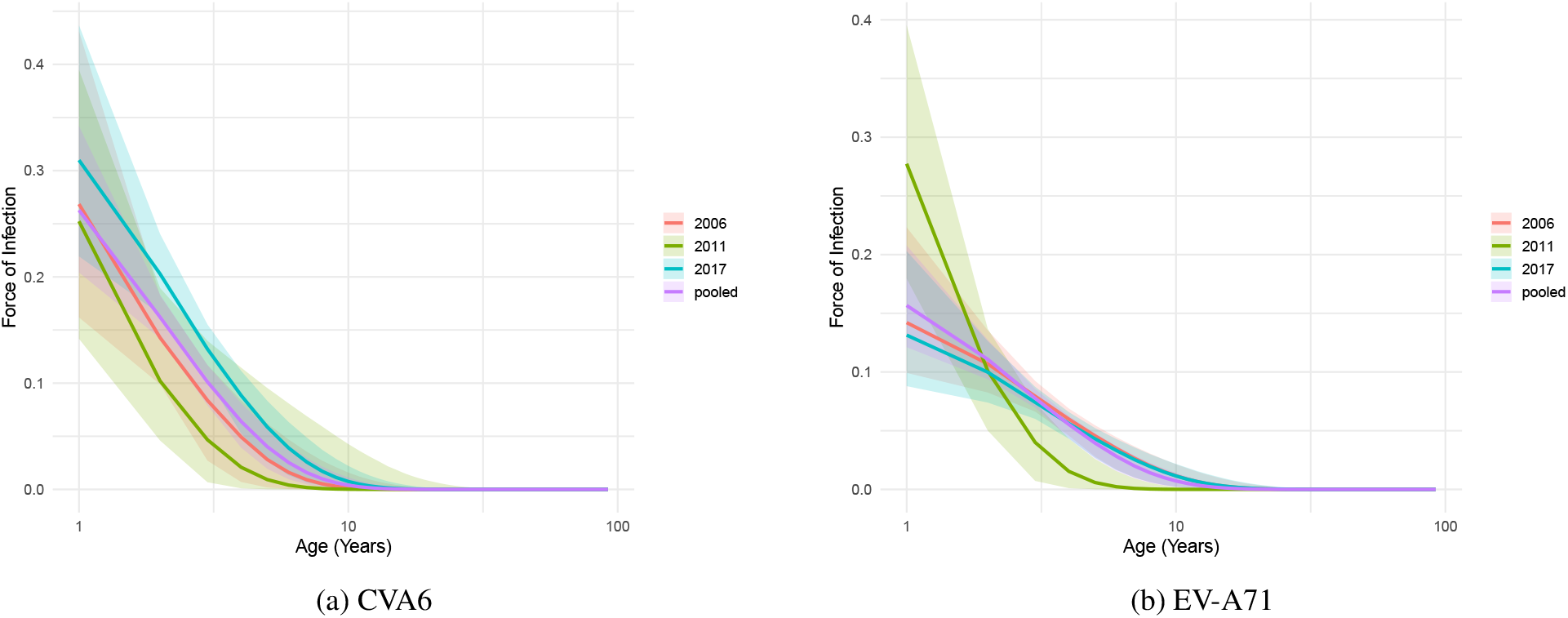
Yearly stratification of the inferred force of infection for CVA6 (L) and EV-A71 (R). Overlaid are the three survey years (2006: red; 2011: green; 2017: blue) and the pooled fits from **Fig. 6** (purple). In all cases the ribbons denote the central 95% posterior credible interval.

**Figure S14:**
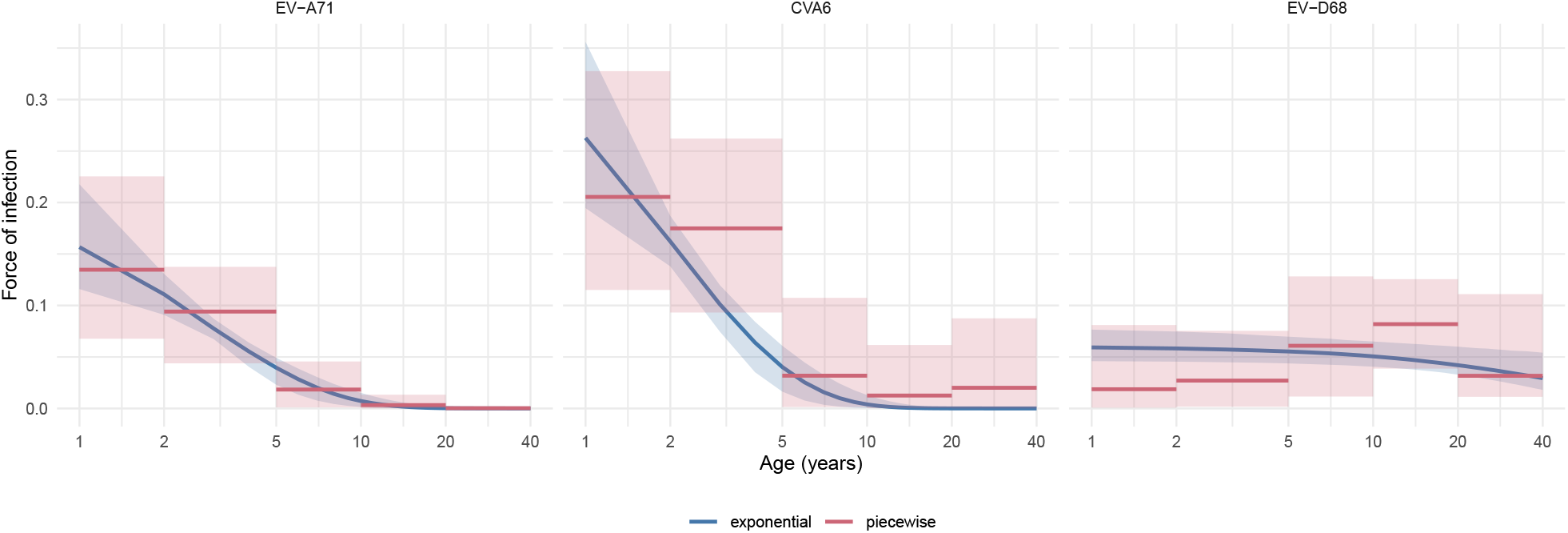
Comparison between inferred exponential (blue) and piecewise (red) force of infection across three enterovirus serotypes. In both cases the ribbon plots denote the central 95% posterior credible intervals. The age brackets for the piecewise force of infection were [1, 2), [2, 5), [5, 10), [10, 20), and [20, 40] years. Approximate LOO-CV did not discriminate between the piecewise and exponential forms for any serotype.

